# Surface curvature and basal hydraulic stress induce spatial bias in cell extrusion

**DOI:** 10.1101/2022.04.01.486717

**Authors:** Cheng-Kuang Huang, Xianbin Yong, David T. She, Chwee Teck Lim

## Abstract

Epithelial cell extrusion is employed in maintaining a healthy epithelium. It remains unclear how environmental conditions specific to various epithelial tissues, such as geometry and fluid osmolarity, affect cell extrusions. We found that, over curved surfaces, epithelial monolayers exhibited higher extrusion rates in concave regions than convex ones. This difference, and overall extrusions, decreased when osmotically induced basal hydraulic stress was reduced by increasing media osmolarity or by culturing monolayers on hydrogels. Mechanistically, basal fluid accumulation antagonized cell-substrate adhesions and the subsequent FAK-Akt survival pathway, leading to apoptotic cell death. Convex surfaces induced cellular forces that acted against osmosis, thereby promoting adhesions and lowering apoptosis. This effect was reversed in concave regions, and together, resulted in a curvature induced spatial bias in cell extrusions.

## INTRODUCTION

Cell extrusion is a coordinated biological process by which epithelial cells are removed from the epithelium without compromising the tissue’s natural barrier function (*1–3*). Typically, in response to external stress, an extrusion disposed cell triggers actomyosin contractions in its neighbors through sphingosine-1-phosphate receptors. The coordinated contraction from its neighbors then squeezes the stressed cell out apically, and concurrently seals the epithelium (*1*). *In vivo*, the stressors that lead to extrusion events include injuries or mutations (*4, 5*), crowding (*6*), and morphogenetic cues (*1*). An often-overlooked fact, however, is that these cell extrusions occur against unique environmental circumstances, such as the characteristic geometries and physical properties of the underlying tissues, and the osmolarity fluctuations in the fluid surrounding them (*2, 7*). It is yet unclear whether these environmental cues can contribute to where and when cells extrude, and how. Interestingly, recent findings have shown that in homeostatic tissue, cell extrusions are by no means spatially random (*8*), as would be expected of a null environmental contribution. Building on the grounds that the biological functions of tissues and organs are often linked to geometric complexities (*9*), as is the case with the eye or an intestinal villus, in this study, we explored whether surface curvature, the defining property of geometric complexity (*10*), can also influence epithelial cell extrusion. Supporting this, curved substrates have been extensively shown to affect morphology (*10–13*), alignment (*14–17*), migration (*18–20*), differentiation (*19–21*) and proliferation (*22*) in single adherent cells.

## RESULTS

### Surface curvature spatially biases epithelial cell extrusions

To explore the effects of surface curvature on epithelial extrusions, repeating wave arrays of hemi-cylindrical valleys and hills with half-period dimensions of approximately 200, 100, and 50 µm (Figure 1A-C) were fabricated using a novel microfabrication procedure (see MATERIALS AND METHODS, Figure 1-figure supplement 1-3, and summary data in Supplementary File 1). Madin-Darby Canine Kidney (MDCK) type II cells were grown to confluency on these substrates for 24 hours, and then live-cell imaged in a Nikon BioStation IM-Q. Figure 1D-F show representative phase contrast images of confluent MDCK monolayers on these waves. The average cell densities prior to imaging, estimated from nuclei staining (Figure 1-figure supplement 4), were 2467 ± 282, 2563 ± 245 and 2509 ± 165 cells/mm^2^ (M ± SD, n = 19, 25, and 19) for 200, 100, and 50 µm waves, respectively.

**Figure 1.**
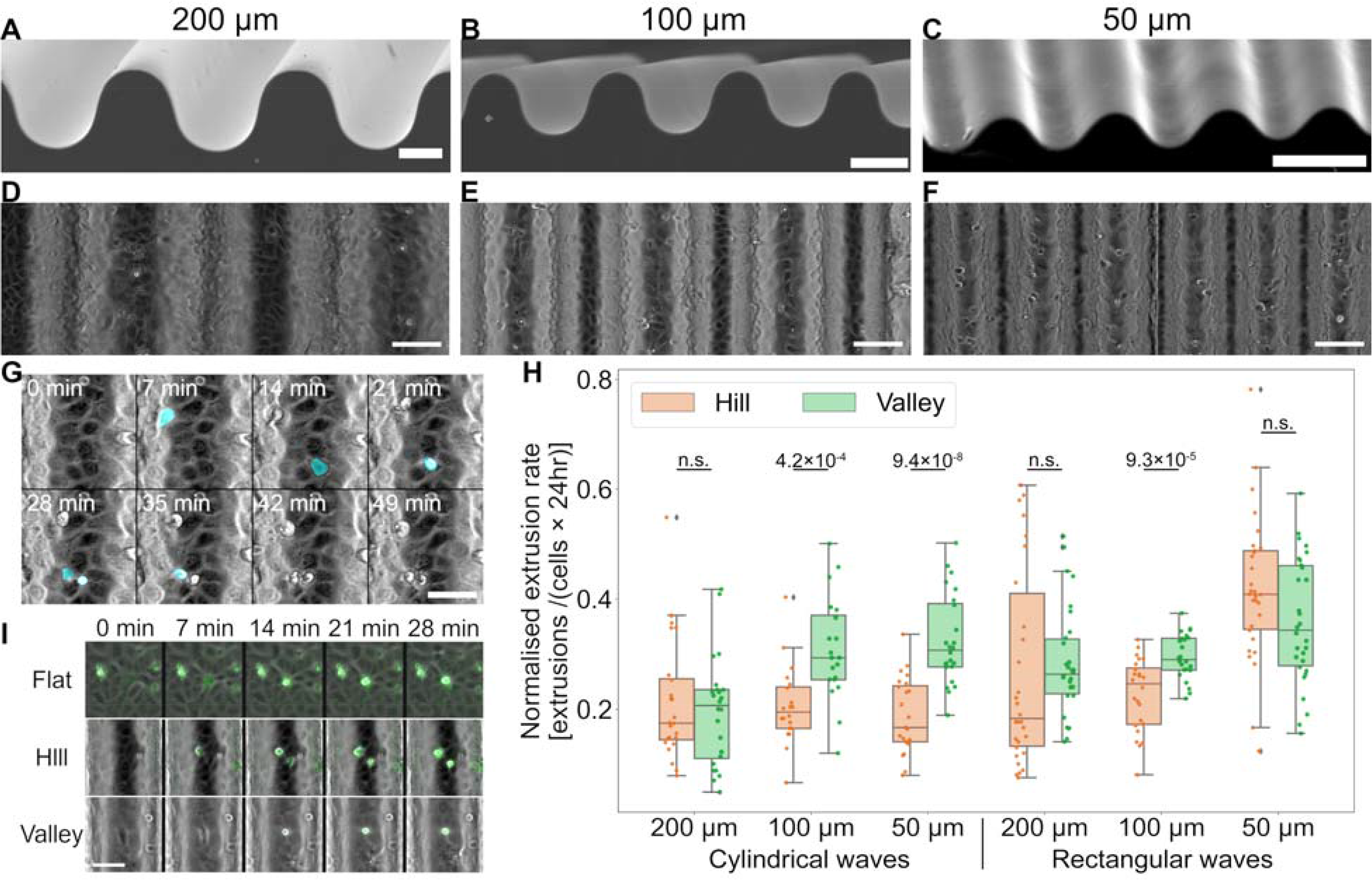
Epithelial monolayers exhibit higher cell extrusion rates in negatively curved valleys of hemi-cylindrical wave substrates. (A-C) SEM images of cylindrical wave structures with half-periods 200, 100 and 50 μm. Scale-bar: 100 μm. (D-F) Phase contrast images of confluent MDCK monolayers on 200, 100 and 50 μm waves, 24 hours after seeding. (G) Time-lapse excerpts demonstrating extrusion event registration (cyan objects) using our trained neural network. (H) Boxplot of extrusion events registered over 24 hours, on 200, 100 and 50 μm cylindrical waves, and on 200, 100 and 50 μm rectangular waves for comparison. Box shows the interquartile range (IQR) while whiskers are 1.5×IQR. Detailed statistics can be found in Supplementary File 2. (I) Multi-channel time-lapse excerpts of cells incubated with activated caspase 3/7 reporter (false green over phase contrast), on flat, hill and valley regions of a 100 μm wave: fluorescence indicate dying cells. Scale-bars (unless otherwise stated): 50 μm.

Qualitative observations of the monolayers over 24 hours revealed more extruded cells in the concave regions (valleys) than convex ones (hills), on the 100 and 50 µm waves (Video1 to 3). To properly quantify this valley-hill extrusion difference, an attention-gated residual U-Net was subsequently trained to differentiate actual extrusion events (false cyan highlights in Figure 1G and Video 4) from floating debris. The architecture of the neural network used in this study is outlined in Figure 1-figure supplement 5A, and a typical training example shown in Figure 1-figure supplement 5B. To validate the performance of the model, a test dataset consisting of 200 positive examples and 100 negative examples were fed into the network and the resulting prediction was obtained from model. The confusion matrix of the model is shown in Figure 1-figure supplement 5C, with examples of false-negative and false-positive events shown in Figure 1-figure supplement 5D. The weighted precision and recall of the model are 0.958 and 0.953 respectively. From this, the normalized extrusion rate was calculated as the number of extrusions per unit area per 24 hours divided by the initial number of cells per unit area (extrusions×cells^-1^×24 hours^-1^).

As shown in Figure 1H (and summary data in Supplementary File 2), a significantly higher mean normalized extrusion rate was found in the valleys compared to hills for monolayers on both the 50 and the 100 µm waves (50 µm valley vs hill: 0.325 ± 0.082 vs 0.187 ± 0.067; 100 µm valley vs hill: 0.308 ± 0.094 vs 0.206 ± 0.071; M ± SD). The valley-hill extrusion rate difference was also higher in the more curved 50 µm wave than the 100 µm wave (0.138 vs 0.103). Meanwhile, no valley-hill extrusion rate difference was found for the 200 µm waves (200 µm valley vs hill: 260.195 ± 0.099 vs 0.213 ± 0.111; M ± SD). These results suggest that epithelial cell extrusions over curved landscapes depended on both the type (concave or convex), and the degree, of underlying surface curvatures.

To test whether elevated valley extrusions were due to potentially less efficient nutrient and mass transfer in those regions, MDCK monolayers were also cultured on rectangular waves of similar dimensions (Figure 1-figure supplement 3D-F, summary data in Supplementary File 1, and Video 5-7). As rectangular corners are essentially highly curved surfaces, extrusion sampling regions were defined to exclude these regions. The cell extrusion rate results shown in Figure 1H (and summary data in Supplementary File 2) revealed no differences between valleys and hills on the 200 and 50 µm rectangular waves, while a small valley-hill extrusion rate difference (Δ = +0.069) was seen on the 100 µm rectangular waves. The lack of similarity between cylindrical and rectangular wave extrusion patterns supports the idea that the valley-hill extrusion rate differences seen in the former were curvature induced.

Epithelial cell extrusions can either be apoptotic, or they can be live in response to extreme crowding (*6*). To test whether the curvature induced extrusion differences were due to surface curvature influencing the relative occurrences of the two extrusion types, we subsequently tracked cell apoptosis. Imaging with the activated caspase 3/7 reporter substrate, CellEvent™, showed that almost all extrusion events, irrespective of location, were accompanied by fluorescence (Figure 1I and Video 8). This indicated that the extrusions were apoptotic, and thus valley-hill extrusion rate differences on the waves were not a result of curvature promoting one extrusion type over the other.

### Osmotically induced basal hydraulic stress is linked to epithelial cell extrusions

Besides valley-hill differential extrusion rates, live-cell imaging beyond 48 hours (Figure 2A and Video 9) typically revealed preferential appearances of fluid-filled domes in the wave valleys. The continuous accumulation of valley domes on the 100 µm waves over this duration is quantified and shown in Figure 2B. As confluent epithelial monolayers function like semipermeable membranes (*23*)—with cellular tight-junctions permitting the passage of water and preventing it for solutes—this separation of the cell monolayer from the substrate indicated an increasing basal hydraulic stress due to osmotic effects. Specifically, over time, water will tend to move into the basolateral compartments with rising sodium concentrations there due to active ion pumping by basolaterally localized Na+, K+-ATPase (*24*).

**Figure 2.**
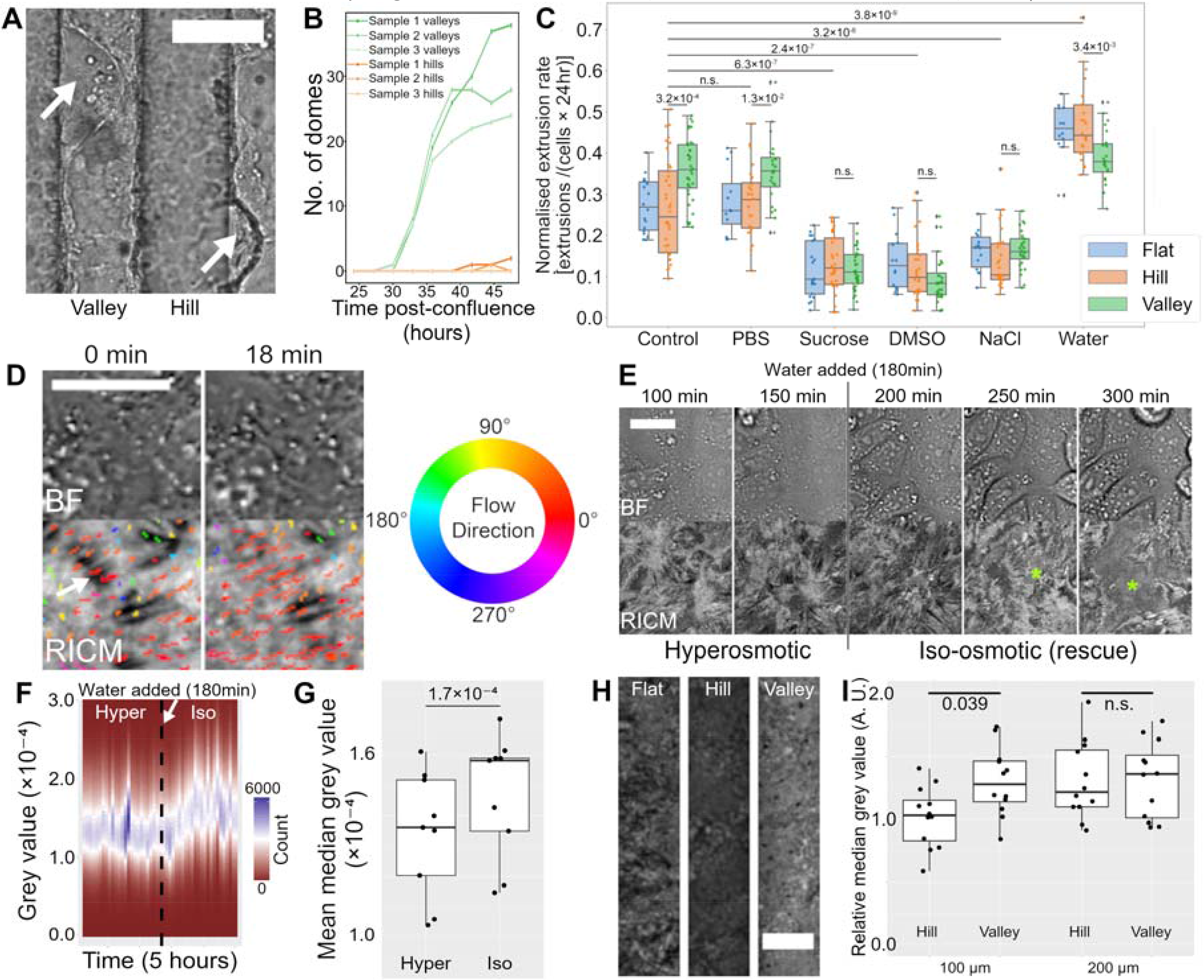
Osmosis induced basal hydraulic stress is linked to cell extrusions. (**A**) Confluent MDCK II monolayers form fluid-filled domes (arrows) in wave valleys when cultured for over 48 hours. Scale-bar: 100 μm. (**B**) scatterplot of valley (green) and hill (orange) region dome-accumulation 24-48 hours into imaging for 3 independent samples (connecting lines are eye-guides). (**C**) Boxplot of extrusions per cell over 24 hours on 100 μm cylindrical waves, with monolayers subjected to osmolarity perturbations that included the addition of 4.1 wt. % sucrose, 1 % DMSO, 0.4 wt. % NaCl, 25 % water, and 25 % PBS. Boxes show the interquartile ranges (IQR) while whiskers represent 1.5×IQR. Detailed statistics can be found in Supplementary File 3. (**D**) Bright-field/RICM time-lapse excepts showing dynamic basal fluid spaces whose motion (direction encoded colored paths) corresponded with the direction of focal adhesion (dark streaks e.g. indicated by white arrow) disassembly. (**E**) Bright-field/RICM time-lapse excepts showing the accumulation of basal fluid (asterisks) when iso-osmolarity was reinstated (at 180 min) in hyper-osmotically pre-conditioned monolayers. (**F**) Plot of the image histograms against time showing an increase in grey values as iso-osmolarity was restored; this indicates a general increase in basal-to-substrate separations with decreasing apical media osmolarity. (**G**) Boxplot of histogram median grey-values from (**F**) averaged separately over the duration of hyper- and iso-osmolarity treatments. Detailed statistics can be found in Supplementary File 4-supplementary file 4a. (**H**) Representative max-projected RICM z-stacks of flat, hill and valley regions (before calibrating against blank substrate intensity differences). (**I**) Boxplot of histogram median grey values from images such as in (**G**) after calibrating against blank sample intensity differences and flat-region medians. Detailed statistics can be found in Supplementary File 4-supplementary file 4b. Scale-bars in (**D)**, (**E)**, (**H**): 20 μm.

We thus asked if basal hydraulic stress could also underly the extrusion differences observed at earlier time points. To test this, and eliminate any non-specific effects from the solutes, we subjected MDCK monolayers grown on 100 µm waves to 3 independent hyper-osmotic conditions—normal media containing either 4.1 wt. % sucrose, 1 % DMSO, or 0.4 wt. % NaCl (Video 10-12). For comparison, a hypo-osmotic condition prepared by diluting normal media with extra 25 % water was also included (Video 13). The measured osmolarities for each condition were 279.3 ± 3.8, 403.6 ± 3.8, 418.6 ± 3.5, 406.3 ± 1.5 and 214.6 ± 4.0 mOsm/L (M ± SD, n = 3) for control, sucrose, DMSO, NaCl and water treatments, respectively.

Strikingly, as shown in Figure 2C (and summary data in Supplementary File 3), by just increasing the osmolarity of the culture media, we were able to significantly reduce the overall cell extrusion rates compared to the control (Control flat: 0.273 ± 0.066, Sucrose flat: 0.115 ± 0.065, DMSO flat: 0.137± 0.070, NaCl flat: 0.162 ± 0.048). In addition, the hyper-osmotic treatments were able to reduce the curvature induced valley-hill extrusion rate differences. Specifically, no difference in extrusion rates between the curvature types (flat, valley, and hill) were detected for all three hyper-osmotic treatments. This also suggests that increasing osmolarities affected valley monolayers more.

Consistent with the effects of hyper-osmolarity, diluting media with extra 25 % water to reduce osmolarity resulted in an increase in overall extrusion rates (Control flat vs water flat: 0.273 ± 0.066 vs 0.457 ± 0.065). Interestingly, valley extrusions did not scale accordingly to maintain a similar valley-hill difference and led to an apparent reversal of the valley-hill extrusion difference (Water valley vs hill: 0.389 ± 0.064 vs 0.470 ± 0.093). This suggests that reduced osmolarities affected hill monolayers more.

Comparison treatments performed by diluting media with 25 % PBS (275.3 ± 0.7 mOsm/L) revealed similar extrusion rates to the control, indicating that the elevated extrusions in our hypo-osmotic treatment were not because of the reduction in nutrient concentration. To further demonstrate that reducing osmolarity by ∼65 mOsm/L does not directly kill the cells, and that the elevated extrusions were contingent upon a full monolayer (i.e., a semi-permeable membrane), we imaged sub-confluent MDCK monolayers subjected to the same hypo-osmotic treatment (Video 14); the cells proliferated normally with negligible cell deaths.

To further support the importance of semi-permeable membrane qualities on cell extrusions, we cultured retina pigment epithelial cells (hTERT RPE-1, ATCC) known to have less developed cell junctions (*25*) on our wave substrates. And, as shown in Video 15, no cell extrusions can be detected from confluent monolayers of these cells over our wave substrates.

As cell adhesions give way to hydraulic stress, the accumulation of fluid is inevitable. To verify the existence of such nascent fluid spaces beneath MDCK monolayers, reflection interference contrast microscopy (RICM) was employed. Relying on the interference of reflected light from the substrate and the cell membrane, the method produces image intensities that are sensitively coupled to basal membrane distances from the substrate (*26–28*). In particular, it has been verified that attached cellular regions (e.g., focal adhesions) appear the darkest, with intensities increasing with basal separation (*26, 27*). Our RICM time-lapse observations (Figure 2D and Video16) indeed revealed numerous continuously moving bright regions (i.e., fluid spaces) under MDCK monolayers on planar substrates. Interestingly, optical flow analysis (*29*) performed on the sequences showed a qualitative correspondence between the directions of the fluid motion (flow paths in Figure 2D) and focal adhesion disassembly (dark streaks fading).

To show that changing osmolarity was sufficient in tuning these basal separations, we tracked variations in the RICM intensities of monolayers grown on planar surfaces and pre-conditioned in hyper-osmotic 4.1 wt. % sucrose media, followed by the addition of water to reinstate iso-osmolarity (Figure 2E and Video17). As osmolarity was lowered back to normal, fluid-filled spaces (asterisks in Figure 2E) begin to develop and eventually span underneath several cells. Moreover, unlike domes (Figure 2A), these nascent basal fluid spaces cannot be inferred from the bright-field images. The minimum grey value aligned histograms of each timeframe plotted against time (Figure 2F) showed the central tendencies of the image histograms shifting toward higher intensities upon the addition of water. By virtue of the physical principal underlying RICM (*28*), this indicated that the basal cell membranes were moving further away from the substrate. Furthermore, the averaged median grey values of the treatment segments (Figure 2G and summary data in Supplementary File 4-supplementary file 4a) showed a ∼9 % intensity increase from hyper-to iso-osmotic treatment segments. This finding strengthens the link between osmotically induced basal hydraulic stress and the osmolarity dependent extrusion rates.

We subsequently examined whether similar basal separation differences exist between valley and hill monolayers on the curved waves. RICM max-projections of MDCK monolayers on the various curvature types of 100 and 200 µm waves showed image contrasts that appeared darkest on the hills and brightest in the valleys (Figure 2H). We subsequently corrected for the drop in reflection intensity in the z direction, due to geometric effects, through imaging blank samples (Figure 2-figure supplement 1). This was then followed by batch normalization against flat regions of each sample to remove cross-sample variation. The results (Fig. 2I and summary data in Supplementary File 4-supplementary file 4b), show that 100 µm-hill relative intensities are significantly different from 100 µm-valleys: hill values were on average ∼23 % lower. In contrast, similar RICM analysis over 200 µm waves did not reveal a significant difference between hill and valley relative intensities. Again, invoking the principal of RICM, these observations suggest that, on strongly curved surfaces, hill monolayers were more closely apposed to the surface as compared to their valley counterparts, paralleling basal separation differences observed between hyper- and iso-osmotic treatments.

To further rule out any cryptic effects in changing osmolarity, we tracked MDCK monolayers cultured on hydrogel (polyacrylamide, PAM) substrates in iso-osmotic medium. Being water and solute permeable materials, hydrogels can buffer against drastic increase in solute concentrations, thereby lowering osmosis induced hydraulic stress. To get a sense of the potential difference in basal solute concentration between the two materials, we can do a quick hand-waving estimation. For monolayers on non-water/solute permeable PDMS of 20×20 mm and using the laser wavelength (640 nm) for RICM as an extreme estimate of basal separation, we should expect ∼0.25 µl of total basal water content. On the other hand, we typically produce our PAM gel slabs using ∼150 µl of precursor solutions. This means that, given similar amounts of solute, PAM gels will lead to monolayer basal osmolarity that is ∼3 orders of magnitude lower than monolayers on PDMS, producing significantly lower osmotic potential.

And to eliminate the possibility that any extrusion differences were due to hydrogels being a much softer material (∼30 kPa (*22*)), MDCK monolayers cultured on soft silicone (CY52-276, Dow Inc., US) of similar stiffness (∼25 kPa (*30*)) were also tracked (Figure 3A and Video18). Figure 3B (and summary data in Supplementary File 5) shows that the extrusion rate differences between the materials were significant. Compared to monolayers on stiff silicone substrates, monolayers on polyacrylamide (PAM) gels produced fewer extrusions over 24 hours (Δ = -0.168). Furthermore, monolayers on soft silicone substrates exhibited higher extrusion rates than their counterparts on stiff silicones (Δ = +0.096). Notably, monolayers on PAM hydrogels typically reached higher cell densities, due to fewer extrusions and therefore more dividing cells (see Video 18 for direct comparison). Similarly, MDCK monolayers in the valley and hill regions of PAM waves also revealed very few extrusions (Video 19), directly paralleling the effects observed when basal hydraulic stress was reduced through hyper-osmotic perturbations (Video 10-12).

**Figure 3.**
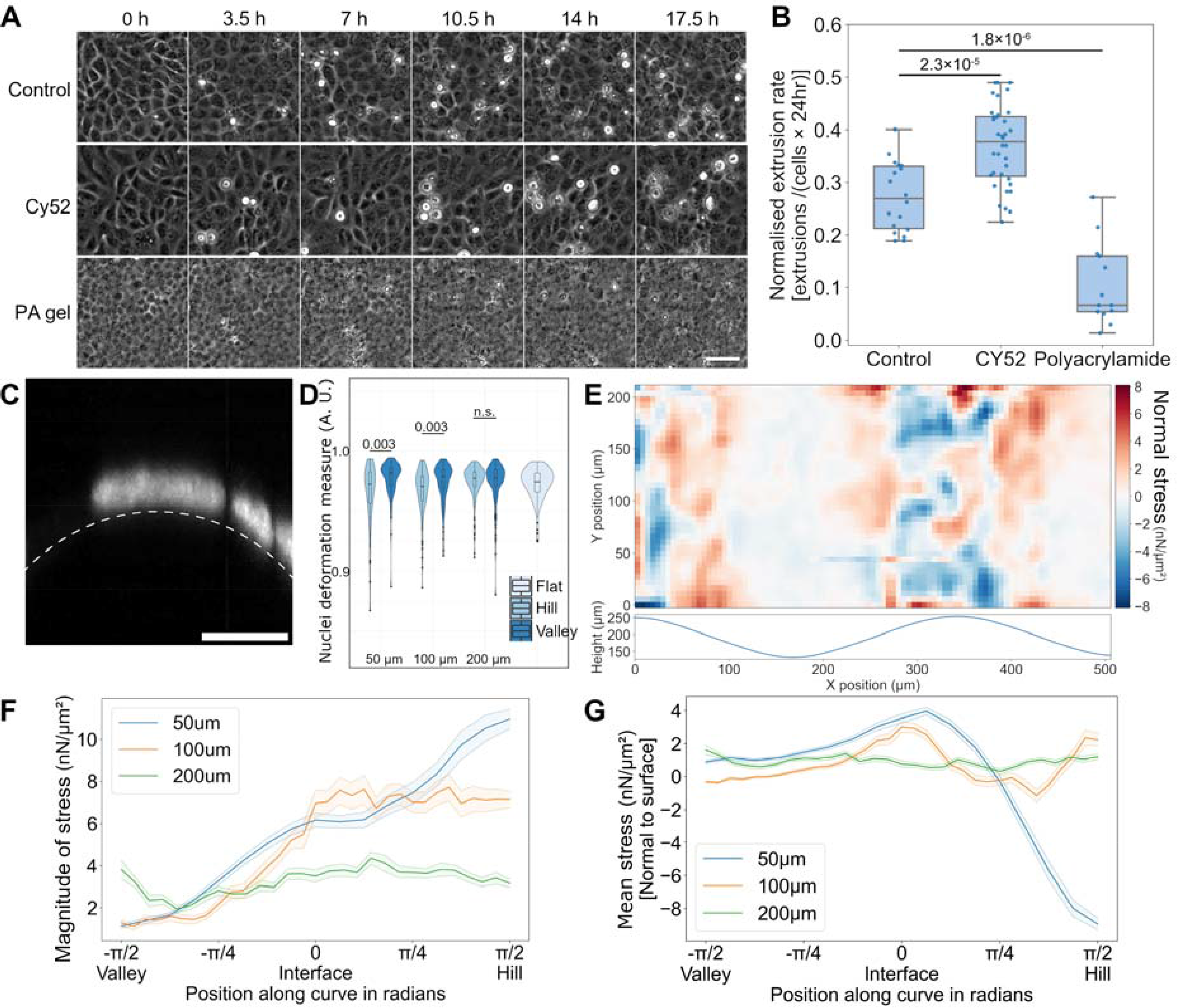
Solute permeable hydrogel substrates reduce epithelial cell extrusions, and surface curvature induces symmetry breaking in collective cellular forces. (**A**) Time-lapse excerpts showing cell extrusion accumulation on stiff PDMS (control), and on soft silicone (CY52-276) and PAM hydrogel of similar stiffness (scale bar: 50 μm). (**B**) Boxplot showing the cell extrusion rates from the 3 substrates in (**A**). Boxes show interquartile ranges (IQR) while whiskers represent 1.5×IQR. Detailed statistics can be found in table S5. (**C**) Fluorescence cross-section showing nuclei deforming against a wave-hill surface (scale bar: 10 μm). (**D**) Violin plot of nuclei deformation measure from cells in valleys, hills, and planar regions, and across the dimension conditions: values closer to 1.0 indicate less deviation from an ellipsoid. (**E**) 3D force-microscopy derived normal stress distribution from a 100 μm wave monolayer: wave profile plotted below. (**F**) Graph showing the bootstrapped magnitude of calculated stresses along the curved profile. (**G**) Mean bootstrapped normal stress vectors along the curved profile. In (**F**) and (**G**), bootstrapping was performed with 10000 re-samples and the 95 % confidence interval is indicated in the shaded region for each respective color.

### Surface curvature induced cell-sheet force asymmetry modulates osmotic effects

While reduced basal separation in hyper-osmotic treatments can be achieved through drawing water from the basal side by osmosis, it is unlikely that neighboring valley and hill regions are subjected to differing osmolarities in the same medium. Instead, judging from geometry, if out-of-plane epithelial forces consistently pointed into the substrates in the hill regions, they would effectively work against osmotically induced basal hydraulic stress and promote adhesion formation and persistence. This could account for the smaller basal-to-substrate separation inferred from RICM.

In support of this, nuclei were readily observed deformed against the surface on the hills (Figure 3C). We quantified this deformation by calculating the ratio segmented-object-volume/fitted-ellipsoid-volume, with values closest to 1.0 indicating least bending. Our quantifications (Figure 3D and summary data in Supplementary File 6) show a statistically significant difference in nuclei deformation measure medians between hill and valley cells on the 50 µm (0.973 vs 0.982) and 100 µm (0.971 vs 0.979) waves. This indicates that cells on the hills tend to have more deformed nuclei compared to cells in the valleys. Meanwhile, no significant difference was found for a similar comparison on 200 µm (0.978 vs 0.978) samples. For reference, the median found for cells pooled from planar regions was 0.975.

We subsequently determined the epithelial force distributions over our wave substrates using a finite element-based 3D force microscopy (see MATERIALS AND METHODS and Figure 3-figure supplement 1). Figure 3E shows a representative monolayer normal-stress distribution on a 100 µm wave, unwrapped and flattened as a 2D projection (algorithm described in MATERIALS AND METHODS). On these waves, the magnitudes of the forces varied consistently over the period, with monolayers on the hills producing larger forces. Interestingly, this effect appeared to be curvature dependent (Figure 3F), with 50 µm waves showing the greatest range of stress magnitudes while the 200 µm structures having an approximately even distribution of stress magnitudes along the surface (50 µm: [1.138, 10.972]; 100 µm: [1.116, 7.725]; and 200 µm: [1.955, 4.356] nN/µm^2^). And as was predicted from nuclei deformation, the directions of the normal forces (Figure 3G) were in general pointing into the substrate on the hills for both the 50 and 100 µm waves. This normal force component decreases as we move away from the hills, flipping to an outward pointing direction at the valley-hill interface. The 200 µm waves did not exhibit any strong curvature dependent force direction patterns, which is consistent with the lack of valley-hill extrusion difference seen earlier. Taken together, this curvature induced force symmetry breaking provided the means for establishing differential basal separation, and the correlated cell extrusion differences, between valley and hill monolayers.

### Basal hydraulic stress antagonizes FAK tyr397 autophosphorylation and its downstream survival signaling

As cell adhesions are intimately related to cell survival (*31*), we hypothesized that basal hydraulic stress may be inducing the apoptotic cell extrusions by antagonizing cell-substrate adhesions. A likely mechanism is through the reduction of focal adhesion kinase (FAK) auto-phosphorylation at tyrosine residue 397 (tyr397), which will ultimately downregulate the downstream Akt survival cascade (*32*). From this, we predicted that the pro-survival effects of hyper-osmolarity would be abolished if FAK inhibitors were introduced together. We tested this by introducing 3 µM of FAKI14 (specific FAK inhibitor at tyr397) to hyper-osmotic 4.1 wt. % sucrose media and imaged for 24 hours (Video 20).

Sucrose was chosen for this and subsequent investigations as mammalian cells in general cannot transport and metabolize polysaccharides (*33*). Without special enzymes to break sucrose down into monosaccharides, such as sucrase found in the gut, the sugars should remain spectators in the culture medium, contributing only to osmotic effects. DMSO, on the other hand, besides changing osmolarity, can also be integrated into cell membrane and pass through cells over time. It has been reported to chronically affect cell membrane properties and gene expressions (*34*). Finally, it is well known that both sodium and chloride ions are readily taken up and transported by cells (*35*). They help to regulate the transmembrane potential, which in turn can affect membrane bound proteins and biochemical reactions within a cell.

As predicted, our results (Figure 4A and summary data in Supplementary File 7) revealed higher extrusion rates when FAKI14 inhibitor was added with sucrose, as compared to the sucrose treatment alone (e.g., FAKI14 flat vs sucrose flat: 0.280 ± 0.073 vs 0.115 ± 0.065), in all three curvature types. In addition, the extrusion rates in FAKI14 treated samples were similar to that in the control for both the flat and valley regions. While the hill region differences in these two conditions were statistically significant, the difference was smaller than the hill region differences between control and sucrose alone. The valley-hill extrusion rate difference was also restored in the FAKI14 treatment (Δ = +0.183) despite higher media osmolarity. Further increasing the FAKI14 concentrations to 6 µM (Figure 4B and Video 21) induced massive cell deaths and led to monolayer disintegration. The pro-survival effects of hyperosmolarity is further supported by the observations that, a lower, 3 µM, FAKI14 concentration was sufficient to elicit similar death response from monolayers in iso-osmotic control media (Video 22).

**Figure 4.**
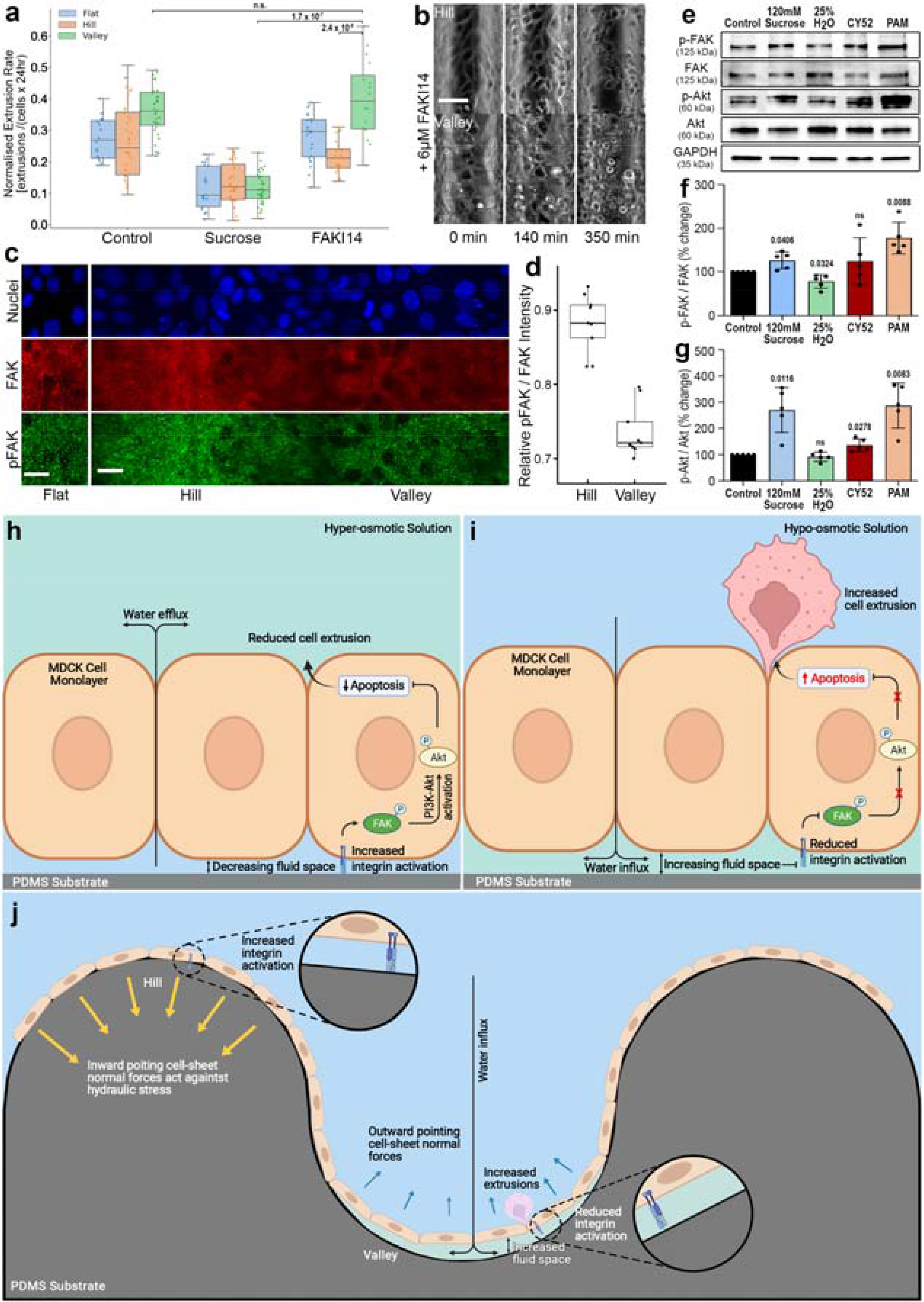
Modulation of basal hydraulic stress through media osmolarity and substrate solute permeability regulate cell extrusion via FAK-Akt pathway. (**A**) Cell extrusion rates on 100 μm waves in normal, sucrose, and sucrose + 3 μM FAKI14 media. (**B**) Adding 6 μM FAKI14 leads to cell death that compromised monolayers. Scale-bar: 50 μm. (**C**) Immunoblots of FAK and Akt proteins in MDCK cells subjected to treatments that lead to varying basal hydraulic stresses for 24 hours; namely, iso-osmotic (control), hyper-(4.1 wt. % sucrose), hypo- (25 % water), iso-osmotic with soft silicone and iso-osmotic with water/solute permeable PAM hydrogel. (**D**) and (**E**) Quantification of the relative expression levels of phosphorylated-FAK (tyr397) as a ratio of p-FAK to total FAK and phosphorylated-Akt (Ser473) as a ratio of p-Akt to total Akt, respectively. GAPDH was used as loading control (M ± SD, n = 5). Statistical analysis can be found in summary data in Supplementary File 8. (**F**) fluorescence images of cell nuclei (false blue), FAK (false red), and p-FAK at tyr397 (false green) on flat, hill and valley; the hill and valley images were unwrapped from 3D stacks. Scale-bar: 20 μm. (**G**) normalized p-FAK/FAK intensities from hill and valley cells. Statistical analysis can be found in summary data in Supplementary File 9. (**H**) and (**I**) Schematic of how osmolarity affect basal hydraulic stress and cell survival. (**J**) Schematic of how curvature induced force differences promote cell survival and death.

To show that modulation of osmolarity affected Akt signaling in a FAK dependent manner, we immunoblotted and quantified the protein expression levels of MDCK monolayers cultured in hyper-osmotic, hypo-osmotic, and iso-osmotic media, and, for comparison, also on soft silicones and PAM hydrogels (Figure 4C-E). Significantly higher levels of phosphorylated-FAK (p-FAK) and phosphorylated-Akt (p-Akt) were observed in media supplemented with sucrose (hyper-osmotic) as compared to those in the iso-osmotic control group. These findings agree with previous reports of higher activated Akt levels in epithelial cells subjected to hyper-osmotic treatment (*36*). Interestingly, FAK activation in the hypo-osmotic condition was not significantly different to the control; this reflects the fact that dead extruded cells cannot be sampled through immunoblotting. Similarly, although the level of p-Akt was reduced under hypo-osmotic treatment, it did not include responses from already extruded cells. On the other hand, cells cultured on PAM hydrogels revealed significantly elevated levels of both p-FAK and p-Akt compared to those on stiff and soft silicones, and the extent of Akt activation was even more prominent (2.5-fold increment) than cells in hyper-osmotic media. Collectively, our findings indicated that monolayers maintained in reduced basal hydraulic stress (hyper-osmotic media and hydrogels) exhibited higher levels of FAK and Akt activation. All else being equal, this will lead to an up-regulation of survival signals that ultimately translates to lower apoptotic cell extrusions.

To show that the same mechanism underlies the valley-hill cell extrusion rate differences over our curved substrates, we visualized p-FAK and FAK on 100 µm waves (Figure 4F). Prior normalizations were applied to the images (see MATERIALS AND METHODS) to compare the relative activation of p-FAK/FAK between hill and valley monolayers (Figure 4G and summary data in Supplementary File 8). In accordance with our observed higher valley extrusion rates, valley cells exhibited ∼20 % lower p-FAK/FAK intensity ratios as compared to hill cells. This predisposes the cells in the valley to a higher likelihood of being extruded as compared to the cells on the hills, resulting in the apparent curvature dependent extrusion rate differences.

## DISCUSSION

Our investigations demonstrate that when cultured on hemi-cylindrical wave substrates, an emergent spatial bias in cell extrusions was elicited in confluent epithelial monolayers. Specifically, MDCK monolayers in valley (concave) regions exhibited significantly higher cell extrusion rates than their counterparts in hill (convex) regions. And this difference was also dependent on the degree of curvature.

Increasing media osmolarity sufficiently reduced this spatial bias and overall cell extrusion numbers, whereas reducing osmolarity increased overall cell extrusions. Imaging planar monolayer basal regions revealed extensive dynamic fluid spaces that responded to osmolarity changes, with lower osmolarity resulting in larger basal fluid spaces and vice versa. As epithelial monolayers functions like semi-permeable membranes, inhibiting solute transport but permitting water, these observations forge a link between the cell-substrate separation induced by osmotically driven basal hydraulic stress and epithelial cell extrusions. On the other hand, on curved substrates without osmolarity perturbations, basal fluid spaces under monolayers in hill regions were found consistently smaller than those in valley regions (as inferred from RICM intensity differences in Figure 2I). With typical extrusion rates higher in the valleys, again, a similar link is established between basal separation and cell extrusions.

We subsequently found that curvature induces out-of-plane cellular forces in hill regions that push cells down on the surface (Figure 4C, G), against osmotically induced basal hydraulic stress and cell-substrate separation, whereas a reversed effect was seen in the valleys. Moreover, the force patterns are in accordance with the curvature induced valley-hill extrusion differences. Specifically, larger valley-hill force differences correlated with larger valley-hill extrusion differences, with 200 µm waves showing no effect in both phenomena. It is worth noting that the protective effects of cellular forces on the hills depend on the strength of apical-basal osmotic gradient. This is evidenced by the fact that hypo-osmotic treatments affected hill monolayers more than valley ones; that is, given sufficient osmotic potential, basal fluid stress can overcome the inward collective cellular forces.

Basal hydraulic stress’ role in epithelial cell extrusions is further supported by the reduced extrusion rates seen, independent of osmolarity perturbations, in monolayers cultured on hydrogel substrates. These types of substrates permit both solute and water and can therefore reduce hikes in solute concentration due to apical-to-basal ion transport by the cells. In our experiments the estimated differences in osmolarity could be as large as 3 orders of magnitude. Ultimately, on hydrogels, the hydraulic stress experienced by the basal membrane would reduce accordingly, promoting adhesion related cell survival. Interestingly, monolayers on the hydrogel substrates typically reach higher cell densities (see Video18 for comparison). We attribute this to the fewer extruded cells and therefore more dividing cells. This also indicates that the cell extrusions seen in our study were not due to cellular crowding.

As basal fluid motion frequently displaced focal adhesions, we hypothesized that the reduced focal adhesion signaling contributed to the cell extrusions observed. Indeed, incorporating focal adhesion kinase inhibitors in high osmolarity treatments re-elevated cell extrusions, indicating the latter’s pro-survival effects were related to cell adhesion formation. Further supporting our hypothesis, immunoblotting analysis revealed higher activation of both FAK and the downstream survival hub protein Akt, in conditions with reduced basal hydraulic stress; that is, monolayers in hyper-osmotic media or on hydrogel substrates. For the apparent elevated levels of AKT seen on soft silicones, we speculate that it is because we cannot immunoblot cells that have died and were inevitably washed out at the start of blotting. Indeed, from the higher extrusion rates seen on these substrates, we could be missing a significant portion of statistics. Specifically, we miss all the cells that would have lowered AKT activation but died, and had we been able to collect those data, lower levels of FAK and AKT may be observed. Alternatively, another explanation could be that, by virtue of survival of the fittest, we might have effectively selected a subpopulation of cells that were able to survive on lower FAK signals, or completely irrespectively of it.

Likewise, on the waves, paralleling the basal separations seen from RICM, higher FAK activation was also observed in monolayers in hill regions compared to their counterparts in valley regions. All else being equal, this will lead to a downregulation of survival signals in valley monolayers and ultimately establishing the apparent spatial bias in extrusion rate differences.

It should be noted that the wave substrates used in our study constitute what is mathematically known as a developable surface. This essentially means that thin structures covering such surfaces experience negligible stretching (*10*). In fact, it is for this property that we can perform unwrapping to reduce our 3D problem into a 2D one. This greatly reduced the effort in our analysis. Structures in the body are rarely this convenient. Based on what we have learned in this study, we can, nevertheless, extrapolate what might be expected of extrusion responses on more complex geometries such as hemispherical surfaces: contiguous (*19*) or discrete (*21*).

For epithelial monolayers in the first scenario, and on poorly solute/water permeable substrates, we should expect to see similar relatively higher extrusion numbers from concave regions compared to convex ones. Moreover, as the surfaces are now curved in both principal directions (likely producing even larger out-of-plane forces), we should see the onset of differential extrusions at larger length-scales. For example, the differences seen on 100 µm hemicylindrical waves might now happen at larger feature sizes for hemispherical waves. Furthermore, as these kinds of surfaces would invariably contain hyperbolic regions (saddle points), we might expect an intermediate response from these locations. If the forces in both principal directions offset each other, the extrusion response may parallel planar regions. Conversely, if one dominates over the other, we may see extrusion responses tending to the dominating curvature (concave of convex).

On the other hand, on curved landscapes with discrete convex or concave regions, we should expect, within curved regions, extrusion behaviors paralleling findings in this study. What would be interesting would be to see what happens at the circumferential rims (or skirt regions) of the features. At these locations we effectively have hyperbolically curved surfaces, and as in the preceding case, we should expect some sort of competing effect between the forces generated from the principal directions. So, for dome skirts, we should see fewer extrusions when the domes are small, and vice versa, when they are larger. Meanwhile, for pit rims, we should see a reversed behavior. The transitioning radial curvature between convex/concave and planar regions would further modulate the effect. Having such a mechanism might be useful in the development of protruding tissues (e.g., villi): the strong curvatures of nascent lumps should favor cell mass accumulation, and once their sizes reach some critical value, epithelial cell extrusions may begin to appear at the roots, offsetting cell division, and eventually halting growth.

The distinctive influences hyperbolic surfaces have on cells is best illustrated in a recent study on epithelial monolayers in bent hydrogel tubes (37). The study found that monolayers on the outside bend tended to detach from the surface whereas those on the inside bend (hyperbolic region) tended to stay well-adhered. The latter can be accounted for if forces from the convex principal direction dominate over the forces from the convex direction. Indeed, the study found an increase in detachment rates with decreasing bend. What is interesting, however, is that the study also reported fewer extrusions occurring from concave regions compared to hyperbolic regions. This is surprising, as the tubes in the study are made of solute and water permeable hydrogel material. And extrapolating from our own observations of epithelial monolayer on hydrogels, we would expect to see minimal differences in the extrusions. A plausible explanation is that the extrusions seen in their study are solely of the canonical crowding effect (*6*). In such a scenario, the detached monolayer on the outside bend could buffer against crowding pressure by buckling. Meanwhile, the monolayer on the inside bend, being attached to the surface, can only regulate crowding pressure by removing cells through extrusion. This phenomenon should be particular to soft friable matrices, such as Matrigel, that separate easily. Using stiffer and covalently bonded ECM should be sufficient to prevent monolayers from detaching, leading to similar crowding extrusions on the outside bend.

In conclusion, recent developments have shown that curved surfaces can direct intestinal epithelial cell differentiation through shaping morphogen gradients (*38*) and influence intracellular YAP/TAZ localization (*39*). Moreover, lumenal hydraulic stress was shown to drive cell patterning in blastocyst development (*40*). Our findings now reveal that surface curvature and hydraulic stress can couple to spatially bias epithelial cell extrusion. Remarkably, basal hydraulic stress alone was sufficient in inducing epithelial cell extrusion, by disrupting cell adhesions and inhibiting downstream Akt survival signals (Figure 4H, I). And an emergent spatial pattern in cell extrusions was elicited when the generation of fluid stress was coupled with curvature induced symmetry breaking of collective cellular forces. Specifically, concave curvatures induced inward out-of-plane forces that acted against basal hydraulic stress, favoring cell-substrate adhesion and survival. Meanwhile, a reversal of this effect occurred on concave surfaces (Figure 4J).

### Ideas and speculations

This emergent multicellular effect has significant implications *in vivo*. The selective removal of cells seen will lead to segregated populations of differentiated cells over time and could well be utilized during development where numerous curved surfaces are formed. The net effect of the collective cellular forces in convex regions is essentially reverse osmosis. It is thus of fundamental interest to explore in future whether such geometric modulation of out-of-plane forces are utilized in a similar way in the body to regulate water and solute transport, in homeostasis and/or development. For example, collective cell contractions on convex surfaces will lead to water exiting the basement membrane, which elevates solute concentrations that might then initiate further cellular responses.

It is also worth noting that, similar to hydrogels in our study, stromal tissues *in vivo* should naturally buffer against osmotically induced hydraulic stress. The capacity to do so will nevertheless vary greatly across different tissue types. This mass transferability can further be impacted by disease. In glaucoma, poor fluid drainage through the trabecular meshwork results in elevated intraocular pressure and ultimately leads to the loss of vision through optic nerve damage (*41*). We therefore foresee our findings to help guide future developments of disease models involving similar dysregulation of water transport.

The realization that osmotically induced basal fluid stress can displace cell-adhesions and, through it, contribute to cell-sheet motion is another important insight. It suggests that basal fluid stress plays a major role in epithelial collective cell migration; the latter drives many biological processes such as wound healing and morphogenesis (*42*). An awareness of fluid stress as a source of epithelial motion in future mathematical models will improve our ability to better understand and explain biological behaviors involving collective cell migration.

Finally, although we have shown that basal hydraulic stress can initiate epithelial cell extrusions, it is also plausible that the positive basal pressure is further utilized in the ultimate physical expulsion of a cell, by the upward momentum the pressure produces. Such an effect could further explain the outstanding question of why cells in some circumstances would extrude basally rather than apically (*1*). Specifically, it is possible that having a higher apical side fluid stress is sufficient in pushing a dying cell into the underlying matrix.

## MATERIALS AND METHODS

### Microfabrication of periodic cylindrical wave structures

The microfabrication of our hemi-cylindrical wave substrates was, in principle, achieved by first manipulating and fusing multiple glass rods to produce an arrayed template. Then, through a series of molding and surface processing, this template was ultimately replicated in a robust monolithic resin format that can be repeatedly molded with silicone elastomer, or other soft biocompatible material, to create curved surfaces for 3D tissue culture. Although the molding of cylindrical structures to create curved substrates has been attempted (*17, 43*), our method can achieve very smooth surfaces. This is in contrast with methods that use metal wires (*43*) which, with their considerable machining defects, may contribute to spurious cellular responses. Furthermore, by selectively removing every other rod from the array, we were able to achieve a contiguous wavy landscape that was symmetric in the dimensions of the concave and convex regions.

Figure 1-figure supplement 1 schematically illustrates in detail the microfabrication procedures involved in producing our cylindrical wave structures. There were essentially 4 major steps in our approach:

**Step 1 (Figure 1-figure supplement 1A).** The first goal was to produce an array of tightly packed uniform glass rods (100 & 200 µm dia., Hilgenberg GmbH, Germany) fixed in place with an UV curable optical adhesive—the Norland Optical Adhesive 73 (NOA73, Norland Products Inc., UK). To facilitate NOA73 lift-off later in the procedure, a glass slide was first pre-treated with Rain-X (800002250, ITW Global Brands, US) according to the manufacturer’s instructions. Then, a thin layer of NOA73 was smeared on to the slide followed by the one-by-one placement of glass rods. The quantity of rods will determine the number of periods produced: for example, 10 rods will produce a pattern with 5 periods. Using a sharp needle under an optical microscope, the rods were pushed tightly together, all the while avoiding trapping debris in between the rods (Figure 1-figure supplement 1A i). Minimal amounts of NOA73 should be used to allow surface tension to hold the rods together; excessive NOA73 will cause the rods to drift apart. To hold the rods in position, the array was then pre-fixed under UV illumination for 1 minutes at 25 % power in a UV-KUB 9” (KLOE, France). After this procedure, two spacers of ∼1 mm thick were placed on either side of the array (Figure 1-figure supplement 1A ii), and additional NOA73 was poured over the array to sufficiently cover it. A second glass slide was used to sandwich the whole construct. This slide was propped up by the spacers and there should be sufficient NOA73 to fill the resulting gap. Again, the NOA73 was cured under UV, this time for 2 minutes at 25 % power. Once the resin has cured, the sandwich construct was pried apart, with separation occurring at the Rain-X treated slide (Figure 1-figure supplement 1A iii).

**Step 2 (Figure 1-figure supplement 1B).** With the NOA73 embedded glass rods, we proceed to remove every second rod in the array to create a wave structure symmetric in its concave and convex curvatures. Residual NOA73 on the top side of the array was first cleared away. Specifically, a sharp needle was pushed along the grooves between the rods, from one edge to the other (Figure 1-figure supplement 1B i), lifting off resin from these regions. Then, with the same needle, every other rod was gently pried from the edge by wedging the needle tip under the rods. To prevent breaking the delicate rods, rather than lifting them off vertically, the needle was run underneath the length of the rods (Figure 1-figure supplement 1B ii). After creating the wave structure, the whole template was gently cleaned using a stiff paint brush, in ethanol followed by water, to remove any NOA73 debris.

**Step 3 (Figure 1-figure supplement 1C).** Due to the finite thickness of the resin between the rods, removing one will unavoidably create a seam along the junctions of neighboring rods (Figure 1-figure supplement 1C i inset). To remove these seams, the glass rod wave template was spin coated with a layer of polystyrene cement (3527C, Testors, US) at 1500 RPM 301 for 10 minutes (WS-400BZ-SNPP/LITE, Laurell Technologies, UK). This extra coat of film will however reduce the concave region’s dimensions while increasing that of the convex ones. Dimensional symmetry was regained by inverting the template followed by a second polystyrene spin coat. To perform the inversion, a two-part silicone elastomer, polydimethylsiloxane (PDMS Sylgard 184, Dow Inc., US), with cross-linker and monomer mixed in a 1:10 weight ratio, was first molded against the original polystyrene-coated template (Figure 1-figure supplement 1C ii). We then created a negative of this PDMS mold by casting against it a second two-part silicone elastomer (Wirosil, BEGO Canada Inc, CA) (Figure 1-figure supplement 1C iii). To prevent fusion of the silicones, the PDMS mold was pre-coated with 1 % bovine serum albumin (BSA, A9647-50G, Sigma-Aldrich) reconstituted in Milli-Q water to act as a de-molding layer. An inverted NOA73 wave template was then derived by UV curing fresh NOA73 sandwiched between the Wirosil silicone mold and a glass slide (Figure 1-figure supplement 1C iv). The resulting inverted template was subsequently spin coated with a second coat of polystyrene, balancing out the dimensional deviations. This substrate constituted our master wave template.

**Step 4 (Figure 1-figure supplement 1D).** The final step of our procedure was to produce multiple resin clones of the master template. These clones, by virtue of parallel fabrication, will facilitate the scale-up production of wave structures for cell culture. For this, we first create a PDMS mold from the master template (Figure 1-figure supplement 1D i). Then, using this mold, NOA73 was repeatedly cast against it, deriving multiple monolithic NOA73 clones on glass slides (Figure 1-figure supplement 1D ii, iii). The clones were subsequently baked at 80 °C for 24 hours before they were used to cast PDMS substrates for cell culture—baking was done to inactivate the adhesion promoter in NOA73 that inhibited PDMS curing. Thereafter, to fabricate wave substrates for cell culture, PDMS was poured onto each clone template and pressed against a large polystyrene culture dish with multiple layers of tape to serve as spacers at either end of the slides (Figure 1-figure supplement 1D iv). When cured, the NOA73 templates were pried from plastic, and the PDMS wave films on the templates were carefully peeled off and attached to 22×22 mm cover-glasses ready for ECM coating (Figure 1-figure supplement 1D v, vi).

The above microfabrication procedure was successfully applied to glass rods with diameters down to 100 µm. Due to the unavailability of commercial glass rods with smaller diameters, we developed an augmentative procedure to accomplish wave structure with half periods smaller than 100 µm. Specifically, drawing inspiration from traditional noodle-pulling, silicone molds were lengthwise stretched to obtain narrower waves. The stretched molds were typically fixed in place on a rigid platform with document clips at either end (Figure 1-figure supplement 2A). NOA73 was then used to replicate stretched molds. To obtain smoother surfaces, this procedure is best performed over several iterations with small stretch increments each time. At each iteration, a new silicone mold was produced from the new narrower wave NOA73 templates to be stretched again (Figure 1-figure supplement 2B). Silicone elastomers with larger elongation at breakage than PDMS, such as Wirosil with a 250 % elongation at break, were employed during the stretching procedures. Furthermore, in our case, a 30 % elongation to the mold was applied at each iteration. This derived periodic wave structures with half periods of ∼50 µm after 4 iterations—based on an assumed 0.5 Poisson ratio.

Using the above methods together, wave structures with half periods of ∼200, ∼100, and ∼50 µm were fabricated (Figure 1A-C). Rectangular waves of similar periodicity to the cylindrical waves were also produced for comparative studies. These structures were fabricated using standard photolithography and soft lithography (*44*).

The surface properties of the PDMS wave substrates, as shown in Figure 1A-C, were characterized using a scanning electron microscope (JEOL JSM 356 6010LV). Meanwhile, to characterize the dimensions of the structures, cylindrical and rectangular, FITC-collagen I from bovine skin (C4361, Sigma-Aldrich) was coated on some of the samples and then imaged on a confocal microscope (see next section). Figure 1-figure supplement 3A-C show orthogonal fluorescence sections of the ∼50, ∼100, and ∼200 µm cylindrical waves, respectively, while Figure 1-figure supplement 3D-F show that of the rectangular waves. Using FIJI and its kappa plugin, the typical dimensions and curvatures (Figure 1-figure supplement 3G, H) as determined from the FITC signals were quantified and are tabulated in summary data in Supplementary File 1.

### Cell culture, staining and imaging

Madin Darby Canine Kidney type II cells (MTOX1300, ECACC collection purchased from Sigma-Aldrich) were cultured in Minimum Essential Medium (MEM, 11090081, Gibco) supplemented with 5 % FBS (10082147, Gibco), 1× Penicillin/streptomycin (15070063, Gibco), 1× GlutaMax, 1× Sodium Pyruvate (11360070, Gibco), and 1× non-essential amino acids (11140050, Gibco). The cell line was maintained at 37 °C in 5 % CO_2_ and sub-cultured every other day. Cells were typically disposed of before 25 passages. hTERT RPE-1 (CRL-4000, ATCC) cells were cultured in accordance with ATCCs recommendations. The MDCK was purchased specifically for this project from ECACC and no batch was used beyond a year of its purchase. The RPEs were kind gifts from professor Rong Li’s lab in MBI and we have thereafter confirmed their authenticity through ATCC services. Both cell lines are tested negative for mycoplasma as a standard requirement and service provided by the wet-lab core of MBI.

For cells to attach to the PDMS wave substrates, an extracellular matrix coating was required on the surface. This was done by first activating the surface under oxygen plasma for 3 minutes. Then, 500 µl of 20 mM acetic acid solution containing 50 µg of collagen-I from bovine skin (C4243, Sigma-Aldrich) was added to the substrate supported on a 22×22 mm cover-glass and left for 2 hours at room temperature. After this, the collagen-I solution was aspirated and the substrate air dried for 1 hour. The coated substrates were then stored at 4 °C and typically used within a week. Prior to experimentation, coated substrates were re-immersed in 1× phosphate buffered saline (PBS, P4417, Sigma-Aldrich) in a 35 mm culture dish and UV sterilized for 15 minutes in a biosafety cabinet.

To study MDCK responses over the curved substrates, cells were first re-suspended in imaging media (MEM without phenol red, above supplements and 2 % serum) followed by seeding on to the substrates at a density of ∼2.5×10^5^ cells/cm^2^. A figure of eight motion was applied to the sample dish to ensure an even distribution of cells. Live-cell microscopy experiments were performed 24 hours after seeding to ensure closure of the monolayer. Otherwise, the monolayers were fixed, at 24, 48 and 72 hours after seeding, for fluorescence imaging.

For nuclei staining, monolayers reaching desired time-points were rinsed in 1× PBS followed by fixing in 4 % formaldehyde (28906, Thermofisher Scientific) for 15 minutes. After two 1× PBS washes, the cells were permeabilized with 0.1 % Triton X-100 (X100, Sigma-Aldrich) for 3 minutes. The cells were then twice washed with 1× PBS, and stained with Hoechst 33342 (H3570, Invitrogen) at 1:1000 dilution. For immunostaining of FAK (610088, BD Transduction Laboratories) and activated FAK (Tyr 397) (AF3398, Affinity Biosciences), cells were first blocked for 30 minutes in 2 % BSA in 1× PBS after permeabilization. The primary antibodies were then diluted 1:400 in the blocking buffer and added to the samples for 2 hours. Afterwards, the samples were rinsed twice in 1× PBS. Secondary antibodies, Alexa Fluor 488 goat anti-rabbit and Alexa Fluor goat anti-mouse (A11008 and A11004, Invitrogen), diluted to 1:400 were then added to the samples and left for another hour. Stained samples were then washed twice in 1× PBS and left in 1× PBS at 4 °C and imaged within a week. The reagents and dilutions for immunoblotting are summarized in Supplementary File 9.

To perform live-cell microscopy, MDCK monolayers grown on substrates attached to cover-glasses were transferred to a stainless-steel cover-glass holder (SC15022, Aireka Scientific, HK) and replaced with fresh image media. The samples were imaged on a Nikon Biostation IM-Q, with the cells maintained at 37 °C and 5 % CO_2_ through the built-in environmental chamber. Phase contrast z-stack images—13 z-planes for all dimensional conditions—were typically acquired every 7 minutes over 24 hours using the internal 10× objective (0.5 NA) and 1.3-megapixel monochrome camera. Furthermore, to obtain a large field of view, a 3-by-5 overlapping image array was tracked for every time-lapse acquisition.

For imaging fixed and stained samples, these were typically propped up by spacers and mounted upside down in 1× PBS on cover-glasses. These samples were then placed in the same stainless-steel cover-glass holder (SC15022, Aireka Scientific, HK) used for live-cell imaging. Fluorescence imaging was then performed on a spinning-disk confocal microscope with a Yokogawa CSU-W1 (Yokogawa Electric, Japan) scanner unit attached to a Nikon Ti2-E inverted microscope (Nikon Instruments Inc., US). Fluorophores were excited through the iLAS laser launcher (Gataca Systems, France) containing 405, 488, 561, 642 nm laser lines. Image z-stacks (z-steps=0.3 µm) were imaged with a 40× water immersion objective (CFI Apo LWD 40XWI λS N.A. 1.15, Nikon) and captured with a sCMOS Camera (Prime 95B 22 mm, Teledyne Photometrics, US). The z-positions were precisely controlled by a piezo stage (PI PIFOC Z-stage, Physik Instrumente, US). The whole microscopy system was controlled by the MetaMorph advanced acquisition software (Molecular Devices, USA).

### Osmolarity perturbation, drug treatment, and apoptosis detection

To create hyper-osmotic conditions, 4.1 wt. % sucrose, 1 % DMSO, and 0.4 wt. % NaCl were added to the imaging media described in the previous section. Meanwhile, a hypo-osmotic condition was created by adding an extra 25 % Milli-Q water. The osmolarity of each condition was measured using the Vapro vapor pressure osmometer 5520 (Wescor Inc., USA). The measured osmolarities were 403.6 ± 3.8, 418.6 ± 3.5 and 406.3 ± 1.5 mOsm/L (M ± SD, n = 3) for the sucrose, DMSO and NaCl hyper-osmotic conditions. Addition of water resulted in a measured osmolarity of 214.6 ± 4.0 mOsm/L. For comparison, the control iso-osmotic media had an osmolarity of 279.3 ± 3.8 mOsm/L.

To test whether inhibiting focal adhesion phosphorylation at tyrosine 397 was sufficient in rescuing cell extrusion rates in MDCK monolayers in hyper-osmotic conditions, image medium containing sucrose was added with 3 µM FAK inhibitor 14 (SML0837-10MG, Sigma-Aldrich) (from 10 mM stock in water) and replaced the iso-osmotic media used prior to imaging. CellEvent™ (R37111, Invitrogen), a caspase 3/7 substrate that produces fluorescence upon caspase 3/7 activation, was employed to visualize the apoptotic proportion of cell extrusions observed in this study. One drop of the reagent was added to 500 µL of medium one hour before live-cell imaging. To minimize phototoxic effects, only four regions, each with 3 z-steps, were imaged for each sample.

To buffer against any increases in hydraulic stress, polyacrylamide hydrogels were fabricated. Specifically, a polyacrylamide precursor solution was prepared by mixing 200 µl of 40 % polyacrylamide solution (Bio-Rad), 200 µl of 2 % bis-acrylamide solution, 580 µl of the MilliQ water and 1.5 µl of TEMED. Next, to 150 µl of the solution, 2 µl of 10 % APS was added and vortexed. This was then added onto the NOA73 molds described in the previous sections before a cover-glasses was placed on top of the mold. The polyacrylamide was allowed to gel for 30 minutes.

To ensure that the polyacrylamide gel binds to the cover-glasses, the cover-glasses were previously treated with a solution containing 2 % trimethoxysilyl propyl methacrylate (Sigma-Aldrich) and 1 % glacial acetic acid in absolute ethanol for 10 minutes. They were then washed 3× in absolute ethanol before being dried in an oven at 80 °C for 2 hours.

The mold was then separated from the polyacrylamide gel/cover-glass structure. This process was done under PBS to ensure that the polyacrylamide gel separates cleanly. The polyacrylamide gel was subsequently washed 3× with PBS before being stored in PBS at 4 °C overnight.

The collagen coating procedure for these hydrogel substrates is described in the 3D force microscopy section.

### Analysis of extrusion rates

Imaging of the various substrates resulted in multiple 3D time-lapse image stacks. To facilitate the downstream analysis of these image stacks, they were unwrapped to obtain a 2D representation of the monolayer. Unwrapping was done by first drawing the side profiles of the wave structures manually. A coordinate map, which detailed the correspondence of the coordinates in 3D space to the 2D plane, was generated from these side profiles. This was then used to unwrap the 3D image stack via spline interpolation.

#### Neural network to detect extrusion events

An attention gated residual U-net was trained to detect extrusion events (Figure 1-figure supplement 5A). The backbone of the encoder and decoder arms of the network followed that laid out previously in (*45*) with the following modifications. The residual block was modified using the ResNet-D block (*46*) with pre-activation (*47*) and the addition of the Squeeze-and-Excitation block (*48*). Furthermore, spatial attention gates were used on the long connections between residual blocks (*49*). Due to the inherent class imbalance between the background pixels and the pixels corresponding to the extrusion events, symmetric unified focal loss was used (*50*). Deep supervision was performed for the various image scales by adding a 1×1 convolution block after each group of residual blocks (*51*). The ground truth was scaled down to the appropriate size and the same loss function as the full-sized ground truth was used. Spatially varying labelling smoothing was applied to the ground truth image as well as every deep-supervision level to mitigate overconfidence in the network (*52*).

Training of the network was performed for a total of 200000 training steps with each step having a batch size of 32. The initial number of filters was set at 64. A cosine learning rate decay, which started from an initial learning rate of 5×10^-4^ and decayed to a minimum of 1×10^-6^, was employed as suggested in (*53*). The training data consisted of input images with 5 frames, 2 before and 2 after the frame of interest, along with a hand-annotated mask of the extrusion events (Figure 1-figure supplement 5B). The intensity of the input images was normalized using percentile normalization. These images were initially 192×192 pixels^2^ in size and were subsequently cropped to the final size of 128×128 pixels^2^ after data augmentation. Both positive and negative examples were annotated to ensure that only extrusion events were being identified by the network. The following data augmentation were used — flips (horizontal and vertical), transpositions and random rotations (multiples of 90°).

As the extrusion process can occur over a few frames, further downstream processing was done to identify the start of the extrusion event. First, a size filter was used to filter out any small objects smaller than 80 pixel^2^. Holes smaller than 80 pixel^2^ were filled as well. Objects with an eccentricity > 0.95 and solidity > 0.5 were also removed. Next, the centroid locations of all the objects were calculated. Objects in different frames were determined to belong to the same extrusion event if the distance between the two centroids were less than 20 pixels and the time difference was less than 5 frames. The first instance of the object in each event was used to represent the position and time of the extrusion. Lastly, only extrusions that occur within the region of interest and the 24-hour timeframe were included in the extrusion rate calculation. The normalized extrusion rate was then calculated by dividing the extrusion rate from each image with the initial cell density for that sample.

#### Estimation of cell density

Cell density was derived using the StarDist network (*54*). Briefly, training images were prepared by hand drawing labels for the cells in representative images using Fiji. The images were also pre-processed using a log-contrast filter to normalize the contrast of the images. The following augmentations were used to augment the training data – random rotation (multiples of 90°), transpositions (horizontal and vertical), flip, motion blur, median blur and Gaussian blur. The training data was also randomly cropped to the final size of 256×256 px before being fed into the network. The default parameters for the StarDist network were used except for the number of epoch and iterations per epoch which were set to 1000 and 100 respectively. Prediction of new images was done using the network weights that provided the smallest loss value during training. By passing the first frame of the time-lapse through the network, the number of cells, and consequently the cell density, were determined for each predefined region of interest (Figure 1-figure supplement 5D). The average initial cell density was then calculated for each sample.

#### Statistical analysis

A total of 3 samples, comprising of 12 images each, were obtained for every condition except for the control which has 4 samples. Statistical analysis of the normalized extrusion rate was performed on these regions of interest using the Python statsmodels module. Two-way ANOVA was performed using the type III sums of squares as well as the HC3 heteroskedasticity-consistent standard error estimators. If the interaction term was significant, the main effects were ignored and only the simple main effects were considered. One way ANOVA of the simple main effects was then performed if multiple comparison were made during post hoc testing. Two-tailed Welch’s unequal variances T-tests were conducted to determine if any of the comparisons were significant. The Benjamini-Hochberg procedure was performed if multiple comparisons were made. Normality of the data was verified via visual inspection of the Q-Q plot and histogram of the residuals. Effect sizes were calculated via bootstrapping with 10000 resamples and presented as the 95% confidence interval.

### Reflection interference contrast microscopy

#### RICM experimental setup

In order to demonstrate the accumulation of water at the basal side of an epithelial monolayer, MDCK monolayers on glass were transferred to stainless steel culture vessels after 24 hours. However, instead of replenishing with iso-osmotic image media described previously, a hyper-osmotic MEM with sucrose was used. Moreover, as the samples were tightly sealed to prevent osmolarity shifts due to evaporation, MEM with Hank’s balanced salt was adopted as the base medium here (11575032, Gibco). After conditioning the cells in hyper-osmotic medium for 4 hours at 37 °C, an air-tight lid was placed on the sample holder. Reflection interference contrast microscopy (RICM) was performed on a Nikon A1R laser scanning confocal microscope operating in reflection mode using 640 laser excitations. The cell-to-substrate attachment was tracked over a 5-hour period with an iso-osmolarity rescue (adding water) performed halfway. The whole system was maintained at 37 °C. The Nikon Perfect Focus System was activated to account for stage drift over time. Furthermore, z-stacks with a 4 µm range and 200 nm intervals were typically acquired to ensure the data always contained the reflection plane of interest.

RICM was also employed to highlight the difference between cell-to-basal attachments between MDCK monolayers on the hills and valleys of the wave substrates. To improve the contrast of RICM, the cells were cultured on wave substrates made from the higher refractive index NOA73 instead of PDMS. After 24 hours, the medium was replenished with iso-osmotic image medium with Hank’s balanced salt, and similarly conditioned for 4 hours prior to imaging. Due to the large vertical span of the structures, the features (hill, valley, and flat) were imaged separately. Typically, z-stacks were acquired with a 20 µm range and 200 nm intervals around the apex of geometries (applicable to hills and valleys only). These values were chosen as RICM would have failed in regions where the surfaces become vertical.

#### RICM data analysis

Our RICM data comprised of z-stacks taken over time and/or over regions. As RICM images are brightest when the plane of reflection is in focus, the reflecting plane at which cells meet the surface was extracted by performing maximum projection in ImageJ (Figure 2D, E, H). Furthermore, being the closest objects to the substrates, focal adhesions should exhibit the lowest intensities (*26, 27*). Utilizing this property, we effectively removed fluctuating backgrounds across time frames by left-shifting the histograms of each frame such that all the minima aligned at zero. This allowed us to register the changing intensities—i.e., the cell-substrate basal separation—over time more accurately (Figure 2F). To derive a quantitative comparison, the medians of the grey values were then averaged over the duration of the treatments (hypo-osmotic or iso-osmotic). Average medians from 3 independent regions and 4 independent samples were compiled (Figure 2G). Normality and equal variance were analyzed using Shapiro-Wilk test and Levene’s test, respectively. Statistical significance was calculated using a two-tailed paired student’s t-test. Effect size was calculated using Cohen’s d.

To make intensity comparisons between cells on the flat, hill and valley regions of 100 and 200 µm waves, rectangular regions of interests were first randomly selected from the max-projected z-stacks of each feature. Again, the histogram minima were left-aligned to zero grey value. The median grey values from these histograms were then compiled—3-4 independent region of interests for each feature from 3-4 independent samples. We then calibrated these values with hill-valley intensity differences obtained separately from blank 100 and 200 µm wave sample RICM images (Figure 2-figure supplement 1A). Finally, to batch normalize the distributions (Figure 2-figure supplement 1B), hill and valley region intensities of each sample were divided by the mean flat-region histogram medians from the same sample (Figure 2I). Normality and equal variance were then analyzed using Shapiro-Wilk test and Levene’s test, respectively. A 1-way ANOVA followed by a post hoc multiple comparison using Benjamini-Hochberg p-value adjustment was applied. The effect sizes were calculated using Cohen’s d.

All data analyses in this section were performed using the statistical programming software, R.

### Quantification of Nuclei deformation

#### Object segmentation

Fluorescent z-stacks of Hoechst-stained nuclei (3 independent sample stacks for each size condition) were first median filtered (1-radius kernel) in FIJI. This was followed by a mean filter (4-radius kernel). The stacks were then thresholded using Huang’s thresholding (not Huang in this study) in FIJI to generate binary image stacks. The 3D object counter plugin was subsequently used to segment and label the objects from the binary stacks. The segmented objects were then imported into the 3D manager plugin (*55*) for curation. Specifically, objects were checked against the original fluorescence z-stack. Merged objects are separated using functions available in the 3D manager, and dividing cells and dead cells are discounted altogether.

#### Nuclei deformation analysis

The volumes of segmented objects and the volumes of best-fit ellipsoids were then measured, and their ratios were exported. The v/v ratios were then imported into R for statistical analysis. The distributions were evidently skewed, thus, a Kruskal-Wallis rank sum test is used as a test for significance. This is followed by post hoc analysis using Wilcoxon rank sum test with Benjamini-Hochberg p-value adjustment (summary data in Supplementary File 6).

### 3D force microscopy

#### Preparation of 3D force microscopy substrate

To study the forces generated by the cells grown on the curved substrates, polyacrylamide substrates with fluorescent beads evenly dispersed throughout the substrate were fabricated (Figure 3-figure supplement 1A, D). Specifically, an aliquot of 10 µl of 0.2 µm fluorescent (580/605) carboxylated microspheres (F8810, Invitrogen) was added to 9.5 ml of Milli-Q water and sonicated for 10 minutes. This solution was then filtered using a 0.45 µm syringe filter (Sartorius). A further 0.5 ml of 500 mM of MES buffer, pH 6.0, was added to the filtered solution. The steps to fabricate the polyacrylamide gel were followed as described in previous sections except for the 580 µl of filtered bead solution being used instead of Milli-Q water.

#### Culturing of cells for 3D force microscopy

To allow for the MDCK cells to attach to the polyacrylamide gel, the surface was coated with collagen-I from bovine skin (C4243, Sigma-Aldrich). This was achieved by functionalizing the gel surface using sulfosuccinimidyl 6-(4’-azido-2’-nitrophenylamino) hexanoate (sulfo-SANPAH). Specifically, 40 µl of 0.2 % acetic acid was added to 2 µl of 10 mg/ml sulfo-SANPAH that was previously dissolved in DMSO. This solution was then added to the polyacrylamide gel surface. A silicone block was used to mechanically agitate the solution to ensure even distribution of the sulfo-SANPAH. UV treatment of the sulfo-SANPAH was done using a UV-KUB 9 (KLOE, France) at 6 % power for 5 minutes. The gel was then washed 3× in 0.2 % acetic acid. This process was repeated a second time with fresh sulfo-SANPAH. Finally, the gel was washed 3× with PBS. 400 µl of PBS containing 50 µg of collagen-I was aspirated onto the gel surface. The collagen was left for 2 hours at room temperature with occasional mixing with a pipette to prevent any collagen gel formation. The gel was then washed 3× with PBS before being stored in PBS. Cell seeding was performed as described previously. After the cell had initially started to attach, the gel was cut using a sharp blade until a small region of interest of approximately 5×1 mm was left (Figure 3-figure supplement 1B). The cells were then allowed to grow for 24 hours until a full monolayer was formed (Figure 3-figure supplement 1C, E).

#### Imaging setup

To image the bead displacements, the gel containing cover-glasses were transferred to stainless steel cell culture vessels and 500 µl of the original media, in which the cell was previously growing in, were added. The vessels were sealed using a custom-made lid to ensure proper humidity control. Imaging of the beads was done using the same imaging system previously used for fixed cells. A z-step size of 0.275 µm was used to ensure that the voxel size was the same for all dimensions. Two sets of fluorescent bead images were captured. The first set consists of a z-stack of beads that was captured with the cell still attached to the gel. The second set consisting of a time-lapse of 5 frames at 15-minute intervals was captured after the cell were removed using 100 µl of sodium dodecyl sulfate (SDS) solution. The SDS solution was prepared by diluting 10 % SDS using the original media to reach a final concentration of 1 %. To minimize any gel deformation, an additional 250 µl of Milli-Q water was added per 1 ml of SDS solution to minimize any gel volume changes due to the addition of SDS. A representative bright-field image of the cells and projected side profile of the fluorescent beads before and after SDS treatment are shown in Figure 3-figure supplement 1F and G respectively.

#### 3D displacement estimation

The displacement of the beads was determined using the two-frame 3D Farneback optical flow method (*56*). We employed 3D Farneback optical flow in our study for its superior computational performance. The method was validated using synthetically generated images from Sample 14 of the Society for Experimental Mechanics DIC challenge. The accuracy of the calculated displacements using the 3D Farneback optical flow was then compared to the provided ground truth displacements. For the highest frequency displacement image pairs, an x-component root-mean-square-error (RMSE) value of 0.0113 was observed. This was lower than the 0.0141 RMSE value for the Augmented Lagrangian Digital Volume Correlation method. This suggested that the 3D Farneback optical flow is capable of accurately calculating the displacement between two bead images.

In this study, a CUDA accelerated implementation was written in Python to speed up the computational time required for displacement estimation. The following parameters for the algorithm were used: 5 iterations, 5 pyramid levels with a scaling factor of 0.5, kernel size of 7 px and box filter size of 31 px. The images were pre-processed by aligning the two sets of images manually before being smoothed with a Gaussian filter of radius 7 px. Validation of the estimated displacement was done by using the estimated displacements to generate a pixel map. This pixel map was used to map the pixels from the bead image without any cells. Qualitative validation was then performed by visually inspecting if there was reasonable alignment between this mapped image and the bead image with cells (Figure 3-figure supplement 1H). Post processing of the estimated vectors was done as follows. First, a suitable region deep in the substrate was chosen and the median x and y displacement from this region was used to eliminate any residual displacement in x and y due to misalignment of the two images. Second, the z displacements along the cross section of the hill were fitted using a best fit line to obtain the z displacement correction function. This function would adjust for any misalignment in z as well as gel shape changes in z because of minor changes to the salt concentration. Last, displacements outside the region of interest were set as undefined.

#### 3D force calculation

The calculation of forces from the displacements were treated as a linear inverse problem. Here, the output (calculated displacements, Figure 3-figure supplement 1I) and the system were known, while the input (forces exerted by the cells) needed to be computed. The system could be obtained from finite element modelling of the gel. The shape of the gel was obtained by subdividing the image into smaller 3D subsets along the axis perpendicular to the feature. Binarization of the image was preformed and the centroid location of the beads were obtained for each subset. The centroid locations were then projected along the axis perpendicular to the feature to obtain a representation of the points on a 2D plane. The alpha shape of the 2D plane was then determined and the outlines for the alpha shapes from each subset were used to calculate the 3D representation of the gel using the “Blend” tool in Ansys Student SpaceClaim 2021 R1 (Ansys Inc, USA). The 3D representation was meshed using Ansys Student Workbench 2021 R1 with the top face having a maximum element edge length of 5.5 µm (Figure 3-figure supplement 1J). The bottom face was set using the zero-displacement boundary condition. To account for any edge effects, the symmetric boundary condition was set for all the faces except for the top and bottom faces. The material properties were specified with a Young’s modulus of 30 kPa and a Poisson’s ratio of 0.457. For each node on the top surface of the mesh, a nodal force of 1×10^-8^ N was applied to the x, y, and z directions sequentially and the corresponding nodal displacement of all the nodes was then recorded. The nodal displacements were then arranged into a matrix describing the response of the system to various nodal forces. The calculated displacements were also mapped to the various nodal positions to obtain the outputs at each node. The inverse problem was then solved via regularization using the Tikhonov method with the regularization parameter determined using the L-curve method (*57*). The surface force per unit area was subsequently calculated by taking the average force of a node and its neighbors and dividing it by the area covered by the nodes (Figure 3-figure supplement 1K).

It should be noted that the force reconstruction method used in this study is implemented from a well-established finite element pipeline to solve inverse problems in engineering and has been repeatedly validated in larger scale engineering contexts (*58*). The novelty and contribution of our article is in its adaptation to reconstruct cellular forces at microscopic scales. To validate the method in our context, we performed simulations of the problem. Specifically, given a finite element model with a predefined curvature, a known force was applied to the surface of the model (Figure 3-figure supplement 2A). The resulting displacements were then calculated from the finite element solution. A 10% random noise is then added to the resulting displacement. The traction force recovery (Figure 3-figure supplement 2B) was then performed using the in-silico noisy displacements. To evaluate the accuracy of the recovery, the cosine similarity along with the mean norm of the force vectors were calculated. A value closer to 1 for both evaluation metrics indicates a more accurate reconstruction of the simulated traction force. The cosine similarity of the recovered traction forces to the original applied force was 0.977 ± 0.056 while the norm of the recovered traction forces as a proportion of the original applied force was 1.016 ± 0.165. As both values are close to 1, this suggested that the traction forces could be satisfactorily recovered using the finite-element based method.

#### Analysis of 3D forces

For each node, the Gram-Schmidt process was used to obtain the orthonormal vectors with the z axis being the normal vector relative to the tangent plane of the node, the y axis being the vector along the long axis of the wave structure and the x axis being the direction that follows along the curvature. Each force was then re-expressed using their respective orthonormal vectors. As cylindrical waves were used, the position of each node can also be expressed as the convex central angle form by the node and a horizontal line cutting across the center of the circle. The bottom of the valley would thus have an angle of -π/2 rad while the top of the hill would have a value of π/2 rad. The interface would have an angle of 0 rad. This facilitated the comparison between the 3 different sizes. The angles were then binned to facilitate the calculation of the mean value as well as the bootstrapped 95% confidence interval. A total of 10000 resamples were used in the calculation of the 95% confidence interval. A total of 6 samples were obtained for the 100 µm waves while 3 samples were obtained for the 50 and 200 µm samples.

### Fluorescence intensity analysis

#### FAK phosphorylation intensity analysis

Due to the 3D nature of our wave substrates, differences in the raw intensities of phosphorylated focal adhesion kinase (p-FAK) fluorescence between hill and valley cells cannot be used directly. To eliminate differences contributed by optical scattering through the material as well as variations in staining, we normalized hill and valley p-FAK and total FAK median grey values against the average median values obtained from 3 random flat regions from the same sample. Then, the p-FAK/FAK ratio, from 3 independent regions and 3 independent samples was calculated and compiled (Figure 4G). Normality and equal variance were determined using Shapiro-Wilk test and Levene’s test, respectively. A two-tailed student’s t-test was used to calculate the significance in the relative activation of p-FAK between cells on the hills and those in the valleys, and the effect size was calculated using Cohen’s d.

All data analysis in this section was performed using the statistical programming software, R.

### Immunoblot Analysis

#### Preparation of MDCK cell lysates

For each experimental condition, the attached MDCK cells were collected by trypsinization (TrypLE Express, Thermo Scientific) supplemented with 50 mM EDTA (Thermo Scientific). The cell pellets were then lysed in radioimmune precipitation assay (RIPA) buffer (Thermo Scientific) containing 25 mM Tris-HCl, pH 7.6, 150 mM NaCl, 1 % NP-40, 1 % sodium deoxycholate, 0.1 % sodium dodecyl sulfate (SDS), along with a protease and phosphatase inhibitor cocktail (Halt™, dilution: 1:100, Thermo Scientific). The lysates were centrifuged at 13,000 rpm for 15 minutes at 4 °C to separate the cell debris from the protein-rich supernatant. The supernatants were then collected into fresh tubes, and protein concentrations were determined by bicinchoninic acid (BCA) assay kit (Thermofisher Scientific) as per manufacturer’s instructions.

#### Immunoblotting

Protein samples were added with equal volume of 2× Laemmli sample buffer (Bio-Rad Laboratories, Hercules, CA, USA) containing 65.8 mM Tris-HCl, pH 6.8, 2.1 % SDS, 26.3 % (w/v) glycerol, 0.01 % bromophenol blue and supplemented with 200 mM dithiothreitol (DTT, Merck). The resultant mixtures were then denatured at 95 °C for 7 minutes and cooled to room temperature. The denatured protein samples (20 µg) were then separated on 4-20 % SDS-polyacrylamide gels using a Tris-glycine running buffer (Bio-Rad) and electroblotted onto nitrocellulose membranes (0.2 µm pore size, Bio-Rad) using a wet Criterion blotter (Bio-Rad) in transfer buffer containing 25 mM Tris base, 150 mM glycine, and 20 % (v/v) methanol for 1 hour at 100 V. The membranes were then incubated in blocking buffer, consists of 5 % (w/v) bovine serum albumin (BSA, Hyclone) in Tris-buffered saline (1× TBST, containing 20 mM Tris-HCl, pH 7.5, 137 mM NaCl and 0.2 % Tween-20) for 1 hour at room temperature to minimize non-specific binding. The membranes were then incubated for 24-48 hours at 4 °C in antibodies targeting the following proteins (dilution factors, incubation duration and manufacturer’s catalogue numbers are tabulated in Supplementary File 8-supplementary file 8c): FAK, phosphorylated-FAK (Tyr397), Pan-AKT, phosphorylated-AKT (Ser473) and GAPDH diluted in blocking buffer. After three washes with 1× TBST, the membranes were incubated in either horseradish peroxidase (HRP)-conjugated goat anti-rabbit IgG (dilution: 1:3000, CST cat# 7074) or horse anti-mouse IgG (dilution factor 1:3000, CST cat# 7076) diluted in blocking buffer for 1 hour at room temperature. The membranes were then subjected to peroxidase-based detection by using enhanced chemiluminescence (ECL) detection reagent (Clarity Max™ Western ECL Substrate, Bio-Rad), and the chemiluminescent signals were visualized by ChemiDoc Touch System (Bio-Rad) and captured by Image Lab software (v 5.2.1). Protein levels were quantified by densitometric analysis using ImageJ software (v 1.53o, National Institute of Health, 755 USA).

#### Statistical analyses

Statistical analyses of all data in this section were performed using 1-sample t-test against the control mean to determine the differences between treatment groups and control group. GraphPad Prism software (v 6.01, GraphPad Software, Inc, La Jolla, CA, USA) was used for these analyses.

## ACKNOWLEDGMENTS

We would like to thank the National University of Singapore (NUS) Mechanobiology Institute for the support, facilities and resources provided during this work. We would also like to thank Dr. Natalia Veronica and Dr. Wan Sia Heng at the Department of Pharmacy, NUS, for assisting in the measurement of the media osmolality, and our colleague Dr. Chii Jou Chan for his suggestions and advice on the paper. X.Y. is supported by the Integrative Sciences and Engineering Programme Scholarship from the NUS Graduate School. We would also like to thank Professor Rong Li for providing the ATCC RPE cells for the revision work.

## DATA AND MATERIALS AVAILABILITY

Data in this study can be obtained as follows:

1. Representative data for the extrusion studies are attached as videos in the main article. Raw extrusion data for these are 24-hour z-stack time-lapses that are 80 GB per file and totals to some 8 TB of data. For this reason, it is difficult to find public repositories that would store these for free over extended times. Nevertheless, the authors would be happy to provide the raw data should the need arise. Specifically, the raw data can be obtained from C. T. Lim at or C.-K. Huang at.

2. Datasets and statistical analysis R scripts for RICM time-lapse, RICM over curved substrates and Nuclei deformation can be obtained from ScholarBank@NUS, a freely accessible repository. Specifically, at https://doi.org/10.25540/XT13 8VRZ.

3. The codes for carrying out finite element assisted 3D force reconstruction, and the companion GPU/CUDA optimized 3D Farneback algorithm, can be obtained again from ScholarBank@NUS at https://doi.org/10.25540/XT13 8VRZ. They are also placed on Github at https://github.com/yongxb/.

4. Similarly, the implementation of the neural network used to detect extrusion events can be found at https://github.com/yongxb/Unet-extrusion-detection and ScholarBank@NUS at https://doi.org/10.25540/XT13-8VRZ.

5. The raw blots and prism files for the immunoblotting study can be obtained from ScholarBank@NUS at https://doi.org/10.25540/XT13-8VRZ.

6. The p-FAK and FAK staining study over the waves and the accompanying analysis can be found in ScholarBank@NUS at https://doi.org/10.25540/XT13-8VRZ.

7. RPE cell-line verification report can be found at ScholarBank@NUS at https://doi.org/10.25540/XT13-8VRZ.

Materials used in this study that are not commercially available, such as the wave substrates used, can also be provided subjected to the terms and conditions of the National University of Singapore.

## List of Supplementary Files

**Supplementary File 1.**

**Table of cylindrical and rectangular wave dimensions.**

**Supplementary File 2.**

**Tables of statistical analysis on varying structure sizes and curvatures. Supplementary file 2a**: 2-way ANOVA analysis to determine if curvature and size of structure have an effect on the normalized extrusion rates. **Supplementary file 2b**: post hoc t-test statistics for the comparisons made with regard to the different sizes of structures.

**Supplementary File 3.**

**Tables of statistical analysis on varying media osmolarities. Supplementary file 3a**: 2-way ANOVA analysis to determine if osmolarity and curvature have an effect on the normalized extrusion rates. **Supplementary file 3b:** 1-way ANOVA of the simple effects on the normalized extrusion rates. **Supplementary file 3c:** post hoc t-test statistics for the comparisons made with regard to the various experimental conditions.

**Supplementary File 4.**

**Tables of statistical analysis of median RICM intensities subjected to osmolarity perturbations and over curvature types. Supplementary file 4a:** two-tailed paired student’s t-test and effect size between the median RICM intensities over hyper- and iso-osmotic treatment timeframes. **Supplementary file 4b:** 1-way ANOVA of normalized median RICM intensities over different curvature and size conditions followed by post hoc analysis with Benjamini-Hochberg (BH) multiple comparison p-value correction.

**Supplementary File 5.**

**Tables of statistical analysis on varying the substrate permeability**. **Supplementary file 5a**: 1-way ANOVA of varying the substrate on the normalized extrusion rates. **Supplementary file 5b:** post hoc t-test statistics for the comparisons made.

**Supplementary File 6.**

**Tables of statistical analysis on the degree of nuclei deformation over wave substrates**. Table shows the medians and IQRs of nuclei deformation measures for each curvature type for 50, 100, 200 µm samples. A Kruskal-Wallis rank sum test is used as a test for significance. This is followed by post hoc analysis using Wilcoxon rank sum test with Benjamini-Hochberg (BH) p-value adjustment.

**Supplementary File 7.**

**Tables of statistical analysis on the addition of FAKI14 inhibitor. Supplementary File 7a:** 2-way ANOVA analysis to determine if addition of FAKI14 and curvature have an effect on the normalized extrusion rates. **Supplementary File 7b:** 1-way ANOVA of the simple effects on the normalized extrusion rates. **Supplementary File 7c:** post hoc t-test statistics for the comparisons made with regard to the various experimental conditions.

**Supplementary File 8.**

**Tables of immunoblotting result statistics and immunoblotting experimental conditions.** Supplementary File 8a: summary statistics and one-sample t-test of p-FAK/FAK expression ratios. Supplementary File 8b summary statistics and one-sample t-test of p-Akt/Akt expression ratios. Supplementary File 8c: reagents and experimental conditions used in the immunoblotting.

**Supplementary File 9.**

**Statistical analysis of p-FAK/FAK fluorescence intensity ratios over hills and valleys.**

## LIST OF VIDEOS

**Video 1.**

24-hour time-lapse of confluent MDCK monolayers on wave substrates with 200 µm half-period dimension.

**Video 2.**

24-hour time-lapse of confluent MDCK monolayers on wave substrates with 100 µm half-period dimension.

**Video 3.**

24-hour time-lapse of confluent MDCK monolayers on wave substrates with 50 µm half-period dimension.

**Video 4.**

Time-lapse showing convolution neural net cell extrusion event registration. Registered events marked out in false cyan.

**Video 5.**

24-hour time-lapse of confluent MDCK monolayers on rectangular wave substrates with 200 µm half-period dimension.

**Video 6.**

24-hour time-lapse of confluent MDCK monolayers on rectangular wave substrates with 100 µm half-period dimension.

**Video 7.**

24-hour time-lapse of confluent MDCK monolayers on rectangular wave substrates with 50 µm half-period dimension.

**Video 8.**

Time-lapse showing caspase 3/7 activation in monolayers on 100 µm wave substrates. Apoptotic cells are accompanied by fluorescence (false green).

**Video 9.**

Valley-hill overlaid time-lapse showing the formation of fluid-filled domes (white arrows) on 100 µm wave substrates when imaged beyond 48 hours. The domes preferentially appear in valleys.

**Video 10.**

24-hour time-lapse of confluent MDCK monolayers on 100 µm wave substrates subjected to hyper-osmotic media containing 4.1 wt. % sucrose.

**Video 11.**

24-hour time-lapse of confluent MDCK monolayers on 100 µm wave substrates subjected to hyper-osmotic media containing 1 % DMSO.

**Video 12.**

24-hour time-lapse of confluent MDCK monolayers on 100 µm wave substrates subjected to hyper-osmotic media containing 0.4 wt. % NaCl.

**Video 13.**

24-hour time-lapse of confluent MDCK monolayers on 100 µm wave substrates subjected to hypo-osmotic media containing 25 % water.

**Video 14.**

24-hour time-lapse of sub-confluent MDCK cells on 100 µm wave substrates subjected to hypo-osmotic media containing 25 % water.

**Video 15.**

24-hour time-lapse of confluent retina pigment epithelial cells (hTERT RPE-1) on wave substrates 100 µm half-period dimension.

**Video 16.**

RICM time-lapse of MDCK monolayers on a planar surface. Dark streaks are locations of adhesions and brighter regions indicate fluids spaces. The time-lapse show numerous fluid spaces that are continuously in motion as indicated by path-lines calculated from optical flow analysis shown in the first panel.

**Video 17.**

RICM time-lapse of MDCK monolayers on planar surface conditioned in hyper-osmotic media before iso-osmolarity is reinstated with the addition of water. The time-lapse demonstrates that monolayer basal fluids are responsive to osmotic perturbations. Several large fluid spaces can be seen towards the end.

**Video 18.**

24-hour time-lapse of confluent MDCK monolayer on planar silicone and polyacrylamide hydrogel substrates. The latter buffers basal hydraulic stress thereby reducing cell extrusions.

**Video 19.**

24-hour time-lapse of confluent MDCK monolayers on hydrogel wave substrates with 100 µm half-period dimension. The buffering of basal hydraulic stress also led to very few extrusions in both valleys and hills. The response parallels that of hyper-osmolarity.

**Video 20.**

24-hour Time-lapse of confluent MDCK monolayers on 100 µm wave substrates subjected to hyper-osmotic media containing 4.1 wt. % sucrose and 3 µM FAK inhibitor 14.

**Video 21.**

24-hour Time-lapse of confluent MDCK monolayers on 100 µm wave substrates subjected to hyper-osmotic media containing 4.1 wt. % sucrose and 6 µM FAK inhibitor 14. Massive cell death leading to compromised monolayers.

**Video 22.**

24-hour Time-lapse of confluent MDCK monolayers on 100 µm wave substrates in control media and 3 µM FAK inhibitor 14. Massive cell death is induced at half the FAKI14 concentration used in media with sucrose.

---

**Figure 1-figure supplement 1.**
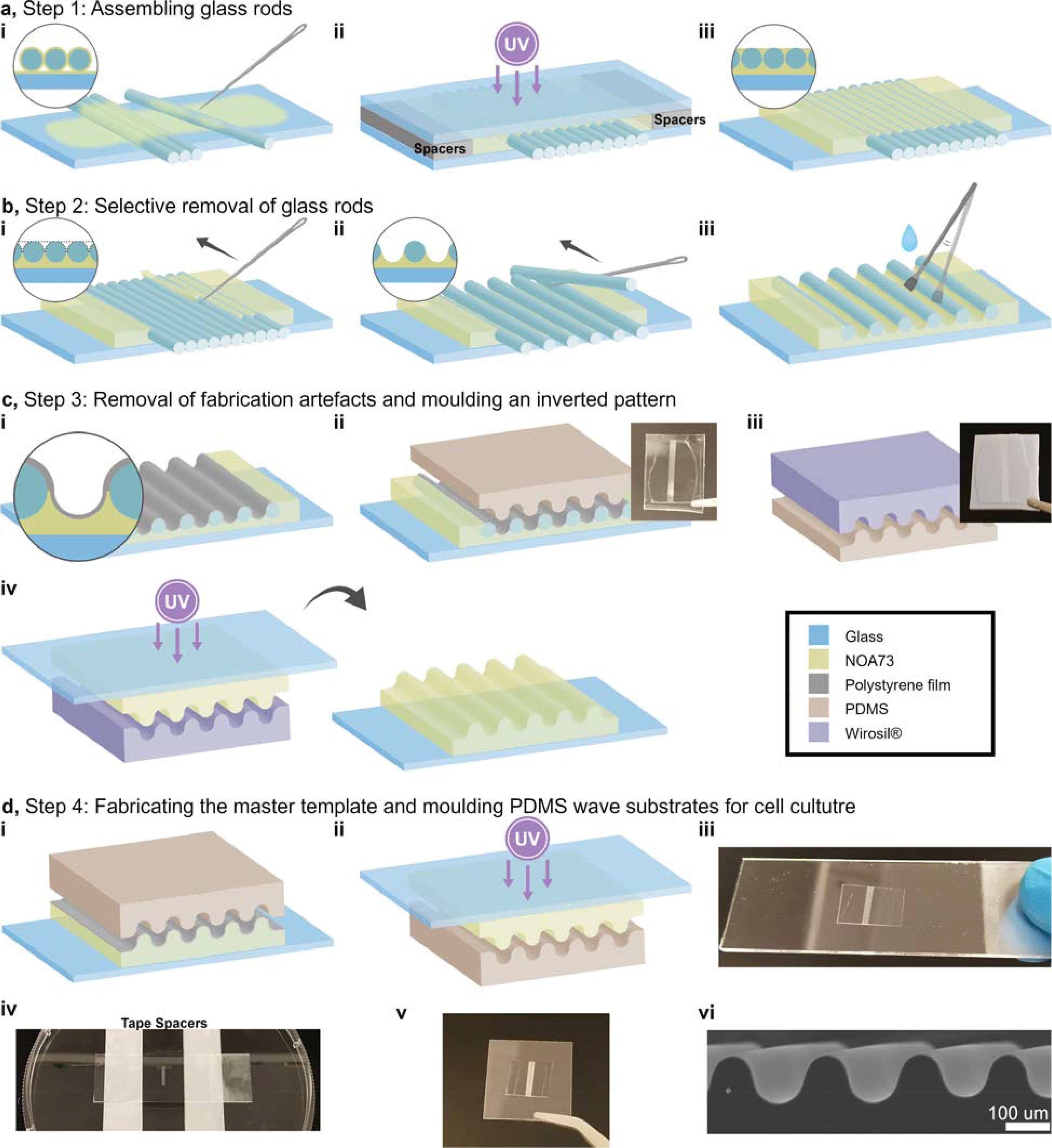
Schematic diagram showing the steps for microfabricating smooth periodic hemi-cylindrical wave substrates using glass rods and iterative molding.

**Figure 1-figure supplement 2.**
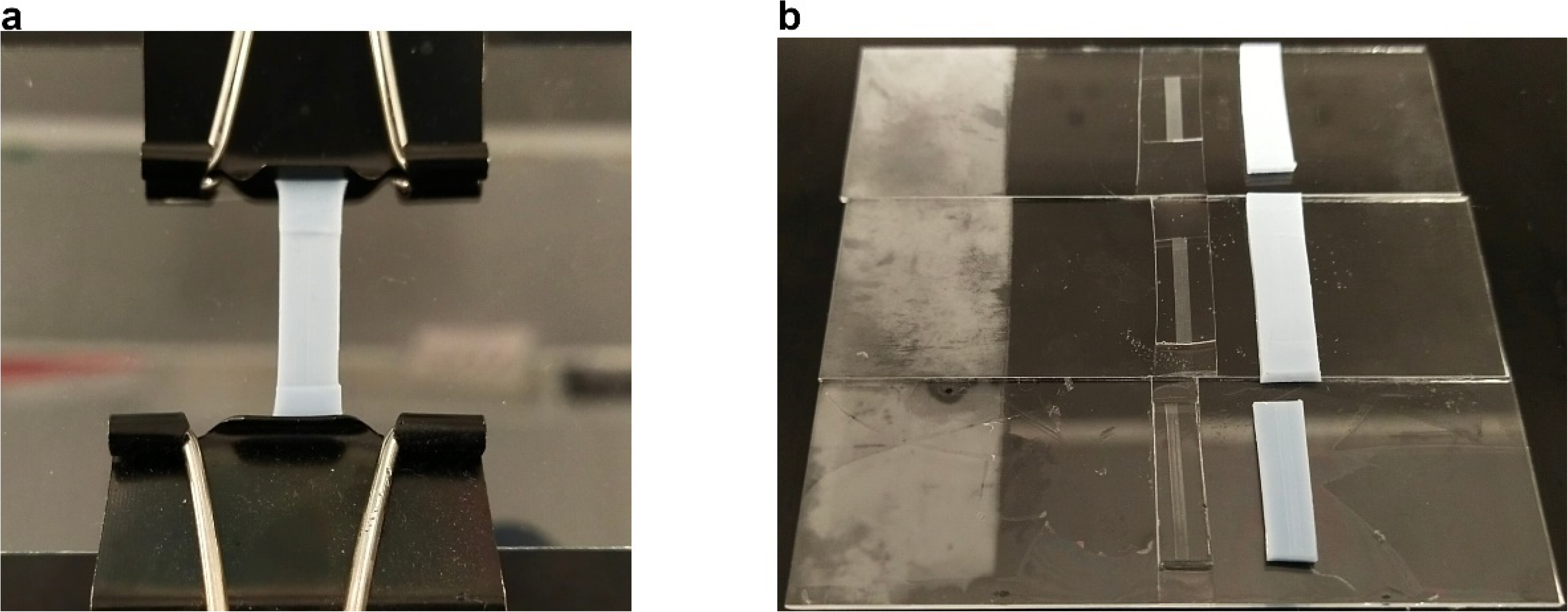
Demonstration of achieving dimensions smaller than commercially available glass rods using an iterative stretching and molding method. **a**, a stretchy silicone with wave pattern was held stretched for optical curable resin casting. **b**, a subsequent stretchy silicone was molded against the new resin template created from a; the whole process was repeated until the desired reduction in dimension was reached.

**Figure 1-figure supplement 3.**
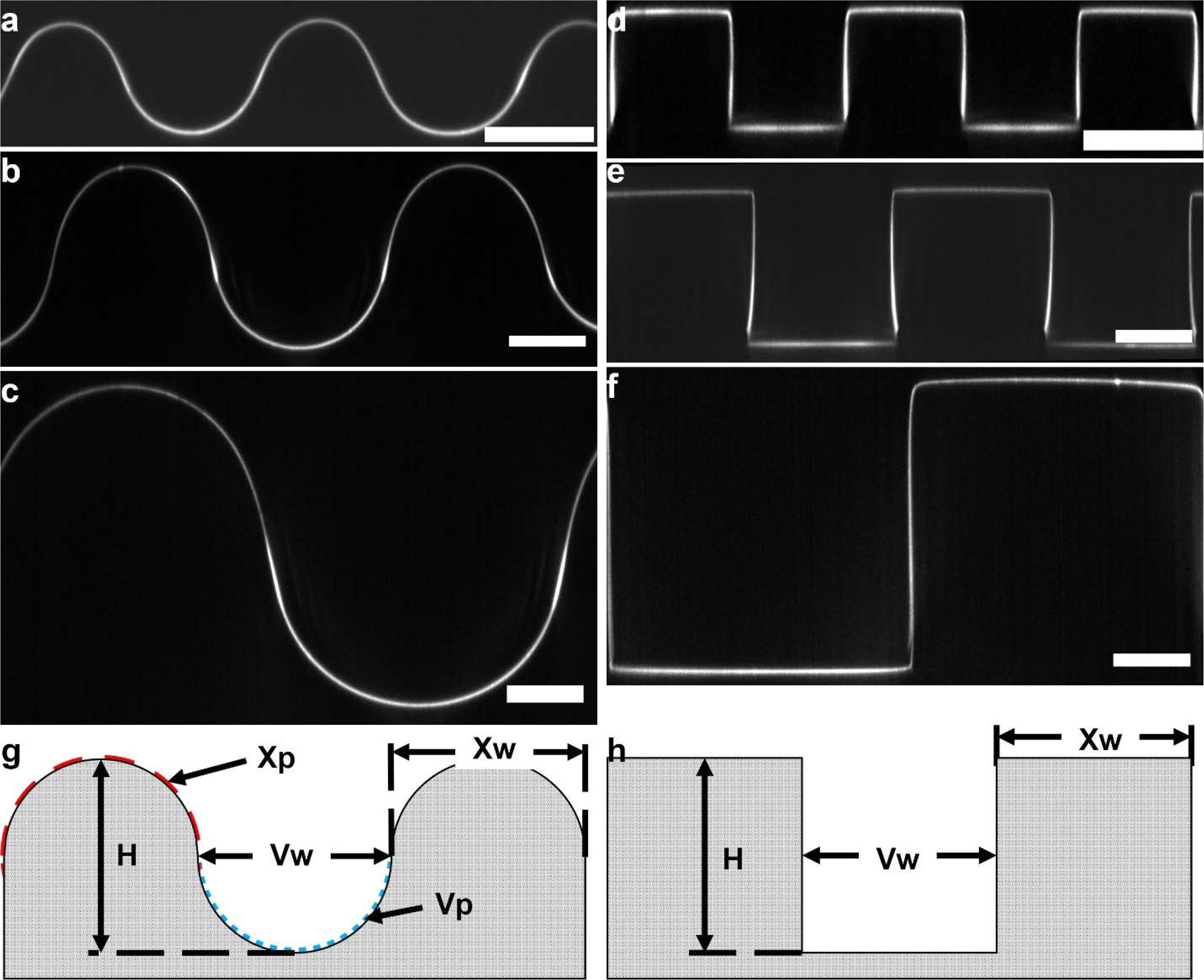
Dimensional characterization of cylindrical and rectangular waves through fluorescent collagen I z-stacks. **a-c**, 50, 100, and 200 µm cylindrical waves. **d-f**, 50, 100, 200 µm rectangular waves. Scale-bars = 50 µm. **g and h**, dimensions measured: curvature (Xp and Vp); width (Xw and Vw); and height (H). Dimension summary data in Supplementary File 1.

**Figure 1-figure supplement 4.**
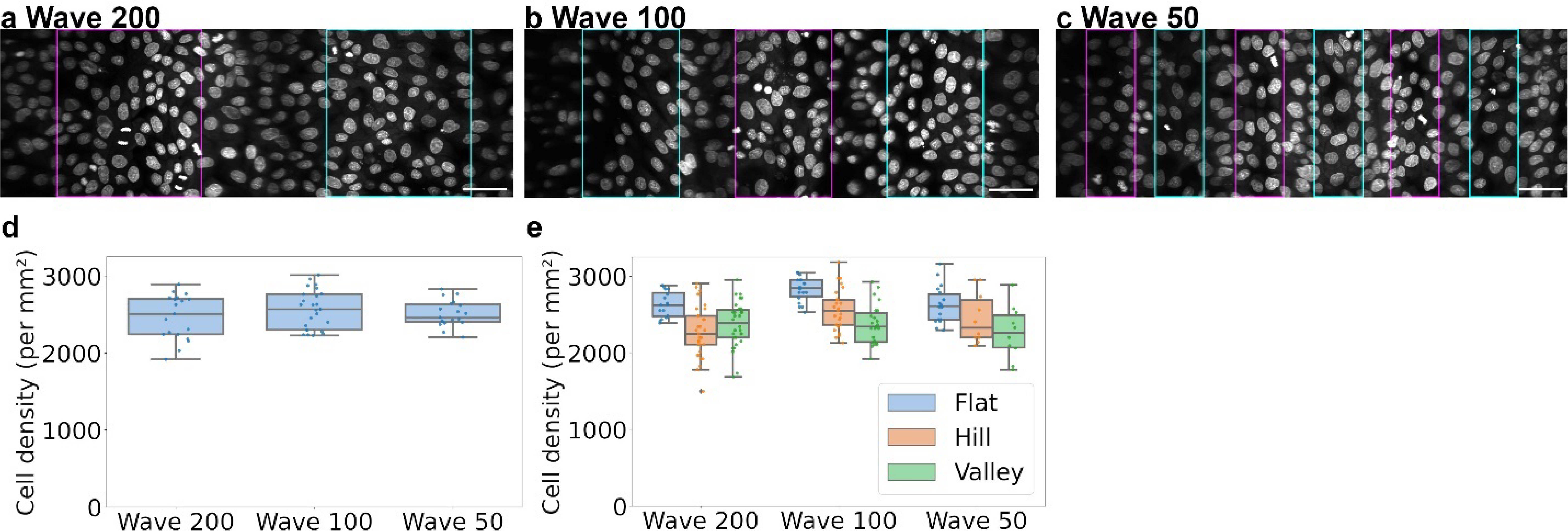
Analysis of cell density 24 hours post seeding. **a-c**, surface unwrapped fluorescent z-stacks of Hoechst stained MDCKs on 200, 100 and 50 µm waves. Hills: cyan boxes and Valleys: magenta boxes. Scale-bars: 50 µm. **d**, boxplot of overall cell densities, determined from nuclei staining, across wave dimensions at the initial time point. **e**, cell densities in **d** plotted according to curvature type.

**Figure 1-figure supplement 5.**
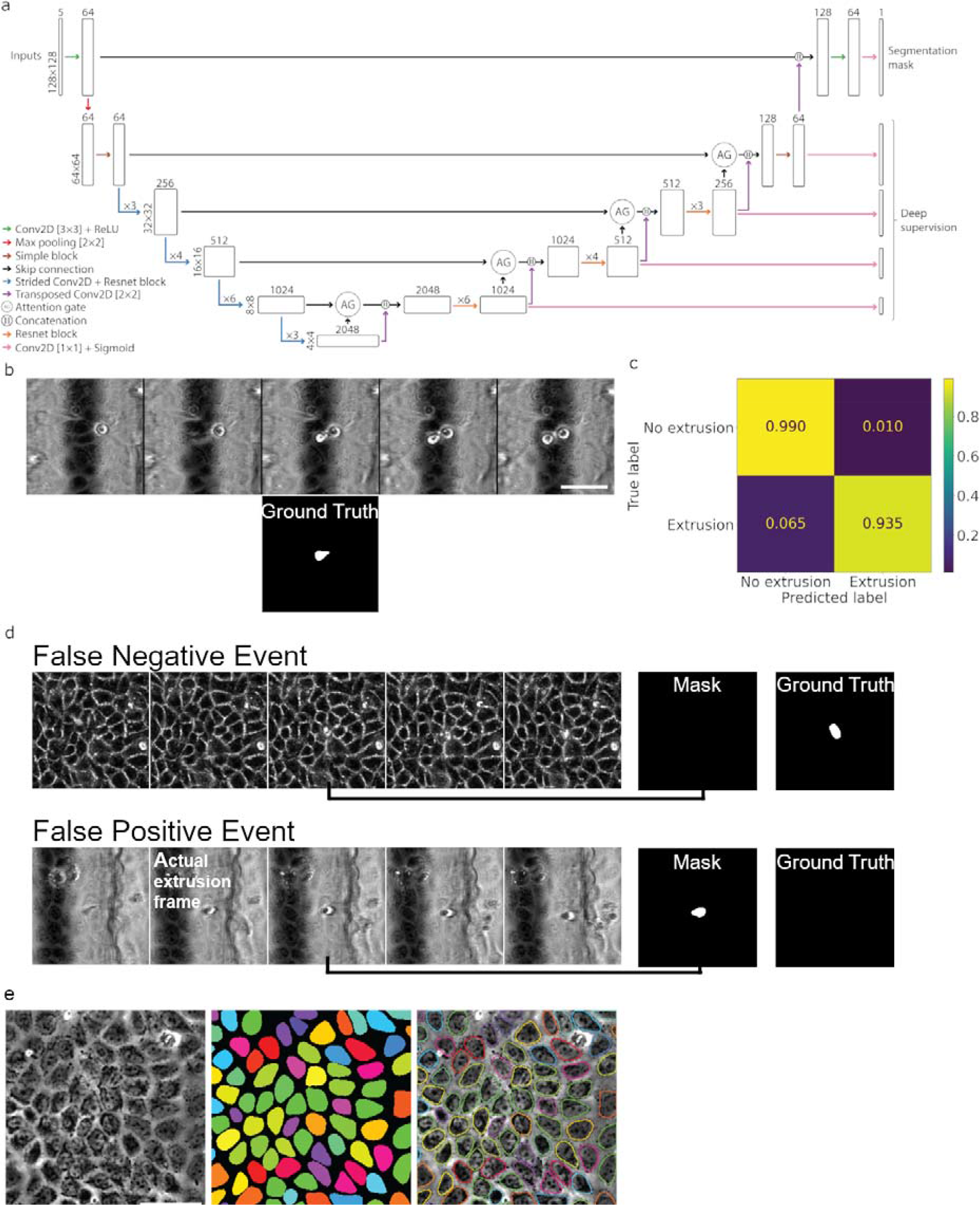
Calculation of the normalized extrusion rate. **a**, architecture of the attention-gated residual U-Net. **b**, example of a manually annotated extrusion event using the sequence of 5 timeframes above and the ground truth binary mask below. **c** confusion matrix of the machine learning network. **d**, examples of false negative and false positive registrations. **e**, Left: bright-field image; Center: prediction from StarDist; Right: overlay of the prediction outline on the bright-field image. Scale-bars: 50 µm.

**Figure 2-figure supplement 1.**
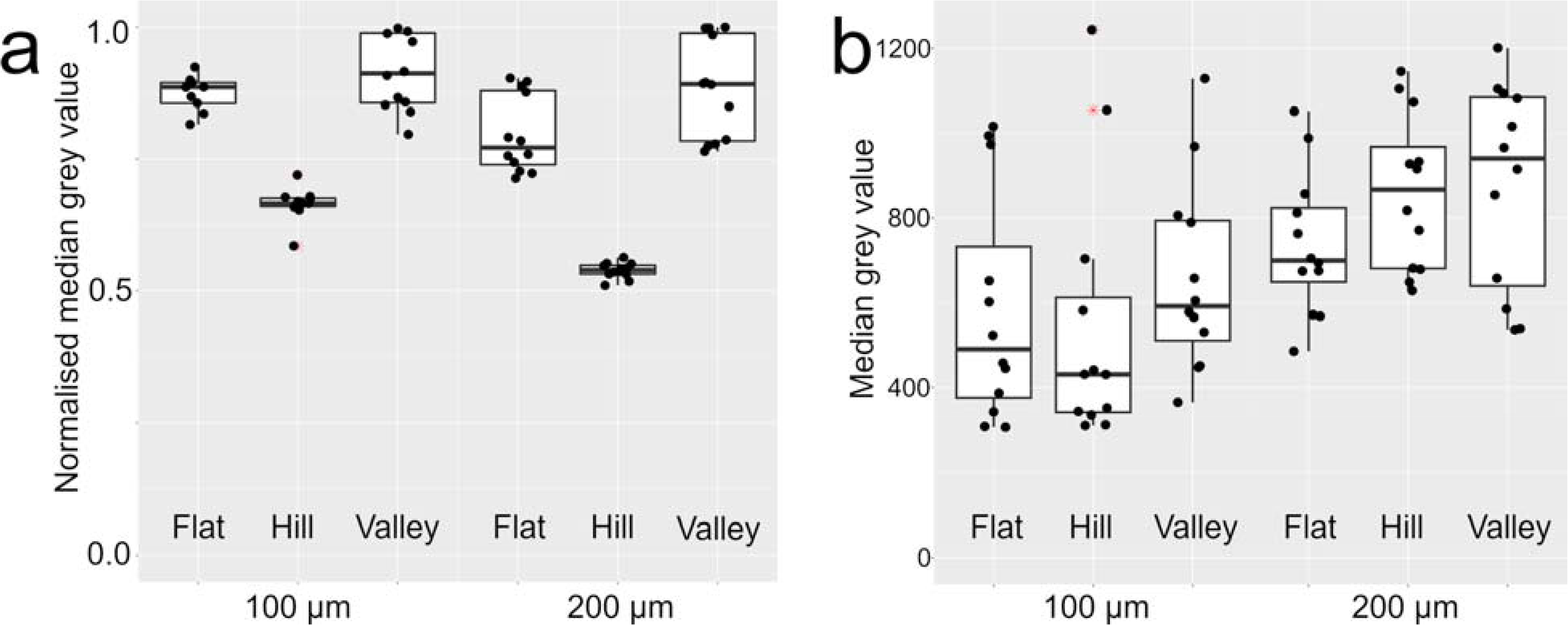
Calibration of RICM intensities against geometric effects. **a**, boxplot showing the reduction of reflected light intensities in hill regions on blank 100 and 200 µm wave substrates. **b**, boxplot of RICM intensities measured from monolayers over 100 and 200 µm wave substrates calibrated against the average reductions inferred from data in **a.** This latter result is then further normalized against flat region signals from each sample batch to remove cross-sample variations, giving the final result shown in main-text Figure 2**I**.

**Figure 3-figure supplement 1.**
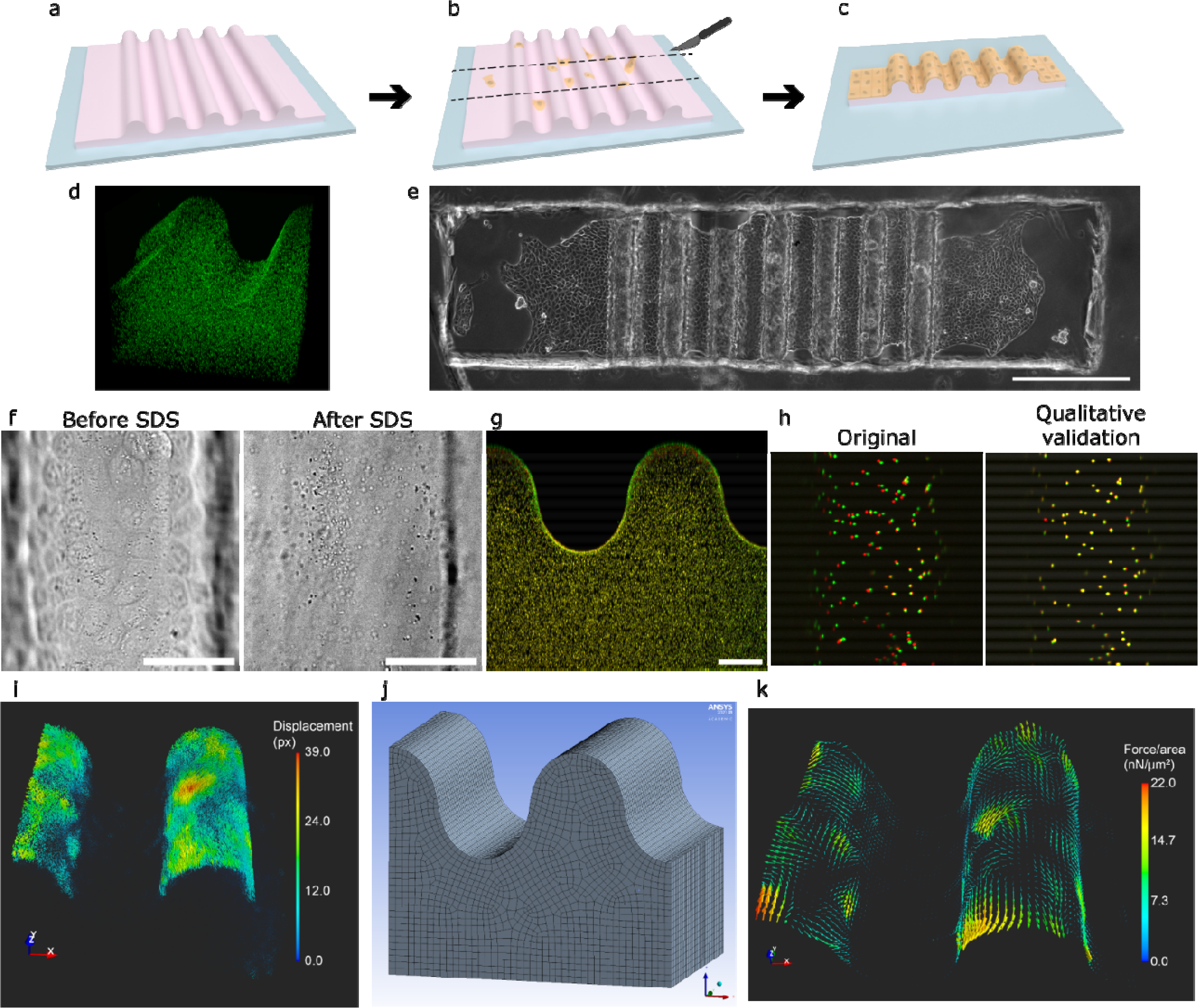
3D force microscopy. **a-c**, schematic for the preparation of the gel as well as the cell seeding process. **d**, 3D-view showing the even distribution of fluorescent beads in the polyacrylamide gel. **e**, bright field image showing the cut get and the monolayer of cells on the curved substrate. Scale-bar: 500 µm. **f**, bright field images showing the removal of cells through the addition of 1 % SDS. **g**, maximum projected side profile of the polyacrylamide gel before (red) and after SDS treatment (green). **h**, Qualitative validation of the calculated displacements. Left: original fluorescent image showing the bead locations with cells (red) and without cells (green). Right: overlay of the mapped image (green) and the bead image with cells (red). **i**, 3D-view of the calculated displacements obtained using the two-frame 3D Farneback optical flow method. **j**, Representative 3D structure and mesh used in the finite element modelling of the polyacrylamide structure. Image used courtesy of ANSYS, Inc. **k**, 3D-view of the average nodal force obtained by solving the inverse problem. Scale-bars: 50 µm unless otherwise stated.

**Figure 3-figure supplement 2.**
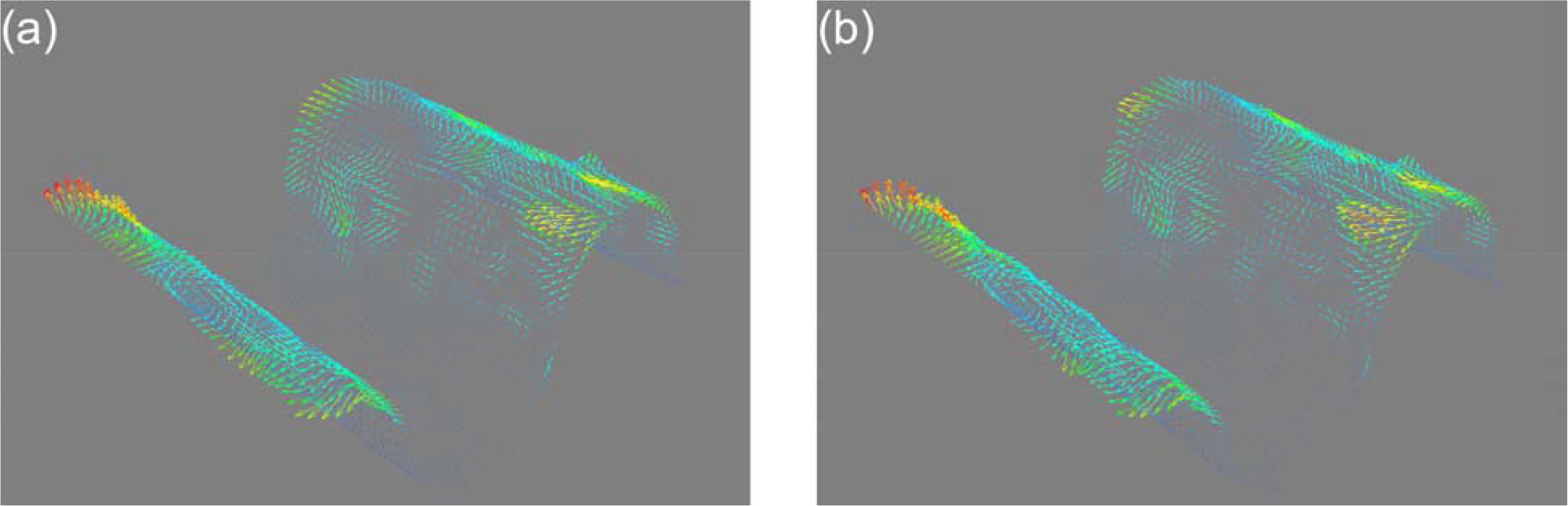
3D force microscopy validation. **a**, shows simulated force field to generate simulated displacements on a 100 µm wave substrate. **b**, shows force field reconstructed from simulated displacements with noise of **a**. See MATERIALS AND METHODS for the quantitative comparison of the two force fields.

