## Supplementary File 1 for "Surface curvature and basal hydraulic stress induce spatial bias in cell extrusion"

| **Substrate type** | **Profile curvature**  **(mean +/- SD μm^-1^)** | **Curvature variance**  **(mean +/- SD μm^-2^)** | **Profile width**  **(mean +/- SD μm)** | **Total height**  **(mean +/- SD μm)** |
| --- | --- | --- | --- | --- |
| 50_cyl  (concave region) | 0.0306 +/- 0.0016 | 0.0163 +/- 0.00010 | 63.6 +/- 3.4 | 53.5 +/- 5.1 |
| 50_cyl  (convex region) | 0.0362 +/- 0.0028 | 0.0218 +/- 0.0019 | 49.5 +/-3.3 |  |
| 100_cyl  (concave region) | 0.0173 +/- 0.0003 | 0.0092 +/0.0008 | 112.1 +/-0.8 | 119.0 +/- 0.4 |
| 100_cyl  (convex region) | 0.0174 +/- 0.0011 | 0.0102 +/- 0.0020 | 104.9 +/-0.6 |  |
| 200_cyl  (concave region) | 0.0107 +/- 0.0005 | 0.0071 +/- 0.0008 | 197.6 +/- 4.6 | 209.3 +/- 3.4 |
| 200_cyl  (convex region) | 0.0105 +/- 0.0008 | 0.0068 +/- 0.0011 | 190.7 +/- 3.4 |  |
| 50_rec  (groove) | - | - | 47.1 +/- 1.2 | 43.0 +/- 1.6 |
| 50_rec  (ridge) | - | - | 47.5 +/- 0.4 |  |
| 100_rec  (groove) | - | - | 97.4 +/- 1.1 | 97.3 +/- 4.3 |
| 100_rec  (ridge) | - | - | 101.8 +/- 2.2 |  |
| 200_rec  (groove) | - | - | 190.8 +/- 1.4 | 187.3 +/- 0.7 |
| 200_rec  (ridge) | - | - | 193.2 +/- 2.4 |  |
| For comparison, curvature of 50, 100, 200 μm circles are 0.04, 0.02, and 0.01 μm^-1^ respectively. | | | | |
