## Supplementary File 2 for "Surface curvature and basal hydraulic stress induce spatial bias in cell extrusion"

**a** 2-way ANOVA analysis of normalized extrusion rates against surface curvature and feature size.

| Source of variation | Sum of squares | DOF | Mean square | F | p-value |
| --- | --- | --- | --- | --- | --- |
| Intercept | 0.958 | 1 | 0.958 | 84.310 | 8.035×10^-18^ |
| Curvature | 0.004 | 1 | 0.004 | 0.330 | 5.664×10^-1^ |
| Size of feature | 0.824 | 5 | 0.165 | 14.508 | 1.061×10^-12^ |
| Interaction | 0.415 | 5 | 0.083 | 7.297 | 1.849×10^-6^ |
| Within | 3.318 | 292 |  |  |  |
| Total | 2.519 | 304 |  |  |  |

**b** Post hoc t-test of valley-hill pairs from different size conditions.

| Group | Sample 1 | Mean | Std | Sample 2 | Mean | Std | Difference | DOF | Welch T | p-value | 95% Cohen D |
| --- | --- | --- | --- | --- | --- | --- | --- | --- | --- | --- | --- |
| Wave 50 | Valley | 0.325 | 0.082 | Hill | 0.187 | 0.067 | 0.138 | 44.2 | 6.374 | 9.372×10^-8^ | [1.144, 2.348] |
| Wave 100 | Valley | 0.308 | 0.094 | Hill | 0.206 | 0.071 | 0.103 | 35.5 | 3.893 | 4.187×10^-4^ | [0.360, 1.857] |
| Wave 200 | Valley | 0.195 | 0.099 | Hill | 0.213 | 0.111 | -0.018 | 45.3 | -0.586 | 5.605×10^-1^ | - |
| Rec 50 | Valley | 0.359 | 0.114 | Hill | 0.418 | 0.132 | -0.059 | 56.8 | -1.852 | 6.922×10^-2^ | - |
| Rec 100 | Valley | 0.295 | 0.041 | Hill | 0.226 | 0.066 | 0.069 | 38.5 | 4.364 | 9.274×10^-5^ | [0.627, 1.739] |
| Rec 200 | Valley | 0.282 | 0.099 | Hill | 0.270 | 0.184 | 0.012 | 44.5 | 0.306 | 7.607×10^-1^ | - |
