## Supplementary File 3 for "Surface curvature and basal hydraulic stress induce spatial bias in cell extrusion"

**a** 2-way ANOVA analysis of normalized extrusion rates against osmolarity and surface curvature conditions.

| Source of variation | Sum of squares | Degrees of freedom | Mean square | F | p-value |
| --- | --- | --- | --- | --- | --- |
| Intercept | 1.630 | 1 | 1.630 | 291.835 | 4.906×10^-50^ |
| Curvature | 0.137 | 2 | 0.069 | 12.265 | 6.663×10^-6^ |
| Osmolarity | 1.405 | 5 | 0.281 | 50.311 | 8.791×10^-41^ |
| Interaction | 0.334 | 10 | 0.033 | 5.981 | 1.701×10^-8^ |
| Within | 2.340 | 419 |  |  |  |
| Total | 5.847 | 437 |  |  |  |

**b** 1-way ANOVA analysis of normalized extrusion rates of the simple effects in Supplementary file 3a.

| Simple effect | Source of variation | Sum of squares | DOF | Mean square | F | p-value |
| --- | --- | --- | --- | --- | --- | --- |
| Hill | Osmolarity | 2.220 | 4 | 0.555 | 70.846 | 4.689×10^-33^ |
|  | Within | 1.128 | 144 |  |  |  |
|  | Total | 3.349 | 148 |  |  |  |
| Valley | Osmolarity | 2.017 | 4 | 0.504 | 138.930 | 2.150×10^-48^ |
|  | Within | 0.523 | 144 |  |  |  |
|  | Total | 2.539 | 148 |  |  |  |
| Flat | Osmolarity | 0.985 | 4 | 0.246 | 60.805 | 1.200×10^-22^ |
|  | Within | 0.300 | 74 |  |  |  |
|  | Total | 1.284 | 78 |  |  |  |
| Control | Curvature | 0.222 | 2 | 0.111 | 12.265 | 2.031×10^-5^ |
|  | Within | 0.786 | 87 |  |  |  |
|  | Total | 1.008 | 89 |  |  |  |
| PBS | Curvature | 0.075 | 2 | 0.037 | 5.446 | 6.846×10^-3^ |
|  | Within | 0.390 | 57 |  |  |  |
|  | Total | 0.465 | 59 |  |  |  |
| Sucrose | Curvature | 0.005 | 2 | 0.003 | 0.740 | 0.480 |
|  | Within | 0.311 | 84 |  |  |  |
|  | Total | 0.317 | 86 |  |  |  |
| DMSO | Curvature | 0.026 | 2 | 0.013 | 2.961 | 0.059 |
|  | Within | 0.290 | 67 |  |  |  |
|  | Total | 0.315 | 69 |  |  |  |
| NaCl | Curvature | 0.007 | 2 | 0.004 | 1.097 | 0.340 |
|  | Within | 0.003 | 67 |  |  |  |
|  | Total | 0.230 | 69 |  |  |  |
| Water | Curvature | 0.091 | 2 | 0.045 | 7.594 | 1.192×10^-3^ |
|  | Within | 0.340 | 57 |  |  |  |
|  | Total | 0.431 | 59 |  |  |  |

**c** Post hoc t-test of the various pairs of experimental conditions.

| Group | Sample 1 | Mean | Std | Sample2 | Mean | Std | Difference | DOF | Welch T | Adj. p-value | 95% Cohen D |
| --- | --- | --- | --- | --- | --- | --- | --- | --- | --- | --- | --- |
| Hill | Control | 0.267 | 0.120 | PBS | 0.283 | 0.087 | -0.016 | 57.6 | -0.602 | 5.496×10^-1^ | - |
|  |  |  |  | Sucrose | 0.134 | 0.067 | 0.133 | 55.8 | 5.758 | 6.296×10^-7^ | [0.961, 1.867] |
|  |  |  |  | DMSO | 0.119 | 0.073 | 0.148 | 59.1 | 6.075 | 2.423×10^-7^ | [1.000, 2.004] |
|  |  |  |  | NaCl | 0.141 | 0.072 | 0.126 | 58.5 | 5.213 | 3.170×10^-6^ | [0.787, 1.808] |
|  |  |  |  | Water | 0.470 | 0.093 | -0.203 | 56.6 | -7.383 | 3.822×10^-9^ | [-1.384, -2.513] |
| Valley | Control | 0.361 | 0.077 | PBS | 0.355 | 0.080 | 0.005 | 48.4 | 0.252 | 8.023×10^-1^ | - |
|  |  |  |  | Sucrose | 0.117 | 0.051 | 0.243 | 60.8 | 15.572 | 1.395×10^-22^ | [3.102, 4.635] |
|  |  |  |  | DMSO | 0.090 | 0.055 | 0.271 | 61.6 | 6.344 | 1.821×10^-23^ | [3.213, 5.112] |
|  |  |  |  | NaCl | 0.165 | 0.045 | 0.196 | 57.7 | 12.694 | 3.210×10^-18^ | [2.46, 3.866] |
|  |  |  |  | Water | 0.389 | 0.064 | -0.029 | 55.2 | -1.558 | 1.248×10^-1^ | - |
| Flat | Control | 0.273 | 0.066 | PBS | 0.282 | 0.080 | -0.009 | 20.5 | -0.310 | 7.599×10^-1^ | - |
|  |  |  |  | Sucrose | 0.115 | 0.065 | 0.158 | 36.0 | 7.504 | 2.895×10^-8^ | [1.846, 3.352] |
|  |  |  |  | DMSO | 0.137 | 0.070 | 0.136 | 27.3 | 5.611 | 5.762×10^-6^ | [1.316, 3.104] |
|  |  |  |  | NaCl | 0.162 | 0.048 | 0.111 | 29.9 | 5.497 | 5.762×10^-6^ | [1.319, 2.780] |
|  |  |  |  | Water | 0.457 | 0.065 | -0.184 | 24.0 | -7.527 | 1.838×10^-7^ | [-1.864, -4.412] |
| Control | Hill | 0.267 | 0.120 | Flat | 0.273 | 0.066 | -0.007 | 51.4 | -0.262 | 7.940×10^-1^ | - |
|  | Valley | 0.361 | 0.077 | Flat | 0.273 | 0.066 | 0.087 | 39.4 | 4.313 | 3.139×10^-4^ | [0.656, 1.863] |
|  | Valley | 0.361 | 0.077 | Hill | 0.267 | 0.120 | 0.094 | 59.9 | 3.945 | 3.174×10^-4^ | [0.458, 1.499] |
| PBS | Hill | 0.283 | 0.087 | Flat | 0.282 | 0.080 | 0.001 | 23.8 | 0.027 | 9.787×10^-1^ | - |
|  | Valley | 0.355 | 0.080 | Flat | 0.282 | 0.080 | 0.073 | 22.0 | 2.591 | 2.498×10^-2^ | [0.243, 1.807] |
|  | Valley | 0.355 | 0.080 | Hill | 0.283 | 0.087 | 0.073 | 45.7 | 3.015 | 1.256×10^-2^ | [0.332, 1.547] |
| Water | Hill | 0.470 | 0.093 | Flat | 0.457 | 0.065 | 0.013 | 30.0 | 0.495 | 6.241×10^-1^ | - |
|  | Valley | 0.389 | 0.064 | Flat | 0.457 | 0.065 | -0.068 | 21.9 | -2.954 | 1.104×10^-2^ | [-0.327, -2.069] |
|  | Valley | 0.389 | 0.064 | Hill | 0.470 | 0.093 | -0.081 | 40.9 | -3.506 | 3.356×10^-3^ | [-0.539, -1.590] |
