## Supplementary File 4 for "Surface curvature and basal hydraulic stress induce spatial bias in cell extrusion"

|  | Statistics | | | Two-tailed Paired t-test | | | | | Cohen D | | |
| --- | --- | --- | --- | --- | --- | --- | --- | --- | --- | --- | --- |
| Condition | N | Mean  (grey value) | SD  (grey value) | t | P | df | 95 % CI | | d | 95 % CI | |
|  |  |  |  |  |  |  | lower | upper |  | lower | upper |
| Hyper-osmotic time segment | 9 | 1.3419.87 | 2050.45 | -6.63 | 0.000165 | 8 | -1666 | -806 | 0.60 (medium) | -0.42 | 1.63 |
| Iso-osmotic time segment | 9 | 14655.78 | 2044.23 |  |  |  |  |  |  |  |  |

(a) Statistics of medium RICM intensities over hyper- and iso-osmotic treatment timeframes.

|  | | Statistics | | | | One-way ANOVA | | | | | | | | | Cohen D | | | | |
| --- | --- | --- | --- | --- | --- | --- | --- | --- | --- | --- | --- | --- | --- | --- | --- | --- | --- | --- | --- |
| Feature size (µm) | Curvature type | N | | Mean  (grey value ratio) | SD  (grey value ratio) |  | | df | Sum sq | Mean sq | | F | P | | d | | 95 % CI | | |
|  |  |  |  |  |  |  |  |  |  |  |  |  |  |  |  |  | lower | | upper |
| 100 | Hill | 12 | | 1.018 | 0.240 | condition | | 3 | 1.021 | 0.340 | | 4.468 | 0.007 | | -1.103  (large) | | -2.012 | | -0.194 |
|  | Valley | 12 | | 1.300 | 0.270 |  |  |  |  |  |  |  |  |  |  |  |  |  |  |
| 200 | Hill | 12 | | 1.307 | 0.307 | residual | | 44 | 3.221 | 0.073 | |  |  |  | -0.021  (negligible) | | -0.868 | | 0.825 |
|  | Valley | 12 | | 1.314 | 0.304 |  |  |  |  |  |  |  |  |  |  |  |  |  |  |
| Post hoc multiple comparison | | | | | | | | | | | | | | | | | | | |
| Pairs | | | n | | | | Statistic | | | | df | | | P | | Adj. P (BH) | | | |
| 100_hill v 100_valley | | | 24 | | | | -2.70 | | | | 21.7 | | | 0.013 | | 0.039 | | * | |
| 100_hill v 200_hill | | | 24 | | | | -3.30 | | | | 19.7 | | | 0.004 | | 0.024 | | * | |
| 100_hill v 200_valley | | | 24 | | | | -2.02 | | | | 21.7 | | | 0.056 | | 0.112 | | n.s. | |
| 100_valley v 200_hill | | | 24 | | | | -0.926 | | | | 20.9 | | | 0.365 | | 0.365 | | n.s. | |
| 100_valley v 200_valley | | | 24 | | | | 0.958 | | | | 20.9 | | | 0.349 | | 0.365 | | n.s. | |
| 200_hill v 100_valley | | | 24 | | | | 1.82 | | | | 18.4 | | | 0.085 | | 0.128 | | n.s. | |

(b) Statistics of normalized median RICM intensities over different curvature types and size.
