## Supplementary File 5 for "Surface curvature and basal hydraulic stress induce spatial bias in cell extrusion"

**a** 1-way ANOVA analysis of normalized extrusion rates on substrate permeability to water and solute.

| Source of variation | Sum of squares | Degrees of freedom | Mean square | F | p-value |
| --- | --- | --- | --- | --- | --- |
| Permeability | 0.585022 | 2 | 0.293 | 53.3434 | 2.330×10^-14^ |
| Within | 0.350947 | 64 |  |  |  |
| Total | 0.936 | 66 |  |  |  |

**b** Post hoc t-test of the various pairs of experimental conditions.

| Sample 1 | Mean | Std | Sample 2 | Mean | Std | Difference | DOF | Welch T | p-value | 95% Cohen D |
| --- | --- | --- | --- | --- | --- | --- | --- | --- | --- | --- |
| Polyacrylamide | 0.105 | 0.078 | Control | 0.273 | 0.065973 | -0.168 | 23.3 | -6.316 | 1.815×10^-6^ | [-0.957, -3.143] |
| CY52 | 0.370 | 0.076 | Control | 0.273 | 0.065973 | 0.096 | 38.9 | 4.801 | 2.349×10^-5^ | [0.618, 1.850] |
