## Supplementary File 6 for "Surface curvature and basal hydraulic stress induce spatial bias in cell extrusion"

|  | | | Statistics | | | | | | Kruskal-Wallis rank sum test | | | | |
| --- | --- | --- | --- | --- | --- | --- | --- | --- | --- | --- | --- | --- | --- |
| Feature size (µm) | Curvature type | | N | | Median | | IQR | | Chi-squared | | df | P | |
| 50 | Hill | | 79 | | 0.973 | | 0.026 | | 35.593 | | 6 | 3.307×10^-6^ | |
|  | Valley | | 83 | | 0.982 | | 0.015 | |  |  |  |  |  |
| 100 | Hill | | 133 | | 0.971 | | 0.020 | |  |  |  |  |  |
|  | Valley | | 118 | | 0.979 | | 0.017 | |  |  |  |  |  |
| 200 | Hill | | 142 | | 0.978 | | 0.013 | |  |  |  |  |  |
|  | Valley | | 125 | | 0.978 | | 0.014 | |  |  |  |  |  |
| Flat | | | 105 | | 0.975 | | 0.016 | |  |  |  |  |  |
| Post hoc analysis: pairwise comparison using Wilcoxon rank sum test with continuity correction (BH p-value adjustment) | | | | | | | | | | | | | |
|  | | 50_hill | | 50_valley | | 100_hill | | 100_valley | | 200_hill | | | 200_valley |
| 50_valley | | **0.003** | | - | | - | | - | | - | | | - |
| 100_hill | | 0.753 | | 3.3×10^-5^ | | - | | - | | - | | | - |
| 100_valley | | 0.061 | | 0.118 | | **0.003** | | - | | - | | | - |
| 200_hill | | 0.052 | | 0.052 | | 0.0013 | | 0.753 | | - | | | - |
| 200_valley | | 0.052 | | 0.114 | | 0.0013 | | 0.956 | | **0.753** | | | - |
| flat | | 0.312 | | 0.008 | | 0.082 | | 0.188 | | 0.184 | | | 0.141 |

Statistics of nuclei deformation measure (segmented volume/fitted-ellipsoid volume).
