## Supplementary File 7 for "Surface curvature and basal hydraulic stress induce spatial bias in cell extrusion"

**a** 2-way ANOVA analysis of normalized extrusion rates against FAK antagonist (FAKI14) and surface curvature conditions.

| Source of variation | Sum of squares | Degrees of freedom | Mean square | F | p-value |
| --- | --- | --- | --- | --- | --- |
| Intercept | 1.901 | 1 | 1.901 | 291.835 | 6.522×10^-43^ |
| Curvature | 0.160 | 2 | 0.080 | 12.265 | 5.664×10^-1^ |
| Treatment | 0.507 | 2 | 0.254 | 38.944 | 2.612×10^-15^ |
| Interaction | 0.303 | 4 | 0.076 | 11.636 | 1.253×10^-8^ |
| Within | 1.511 | 232 |  |  |  |
| Total | 4.383 | 241 |  |  |  |

**b** 1-way ANOVA analysis of normalized extrusion rates of the simple effects in Supplementary file 7a.

| Simple effect | Source of variation | Sum of squares | Degrees of freedom | Mean square | F | p-value |
| --- | --- | --- | --- | --- | --- | --- |
| Hill | Osmolarity | 0.340 | 2 | 0.170 | 21.708 | 2.026×10^-8^ |
|  | Within | 0.705 | 90 |  |  |  |
|  | Total | 1.045 | 92 |  |  |  |
| Valley | Osmolarity | 2.017 | 4 | 0.504 | 138.930 | 2.150×10^-48^ |
|  | Within | 0.523 | 144 |  |  |  |
|  | Total | 2.539 | 148 |  |  |  |
| Flat | Osmolarity | 0.985 | 4 | 0.246 | 60.805 | 1.200×10^-22^ |
|  | Within | 0.300 | 74 |  |  |  |
|  | Total | 1.284 | 78 |  |  |  |
| FAKI14 | Curvature | 0.242 | 2 | 0.121 | 17.806 | 8.113×10^-7^ |
|  | Within | 0.414 | 61 |  |  |  |
|  | Total | 0.655 | 63 |  |  |  |

**c** Post hoc t-test of the various pairs of experimental conditions.

| Group | Sample 1 | Mean | Std | Sample 2 | Mean | Std | Difference | DOF | Welch T | Adj. p-value | 95% Cohen D |
| --- | --- | --- | --- | --- | --- | --- | --- | --- | --- | --- | --- |
| Hill | FAKI14 | 0.217 | 0.051 | Sucrose | 0.134 | 0.067 | 0.083 | 54.9 | 5.335 | 3.724 | [0.896, 1.983] |
|  |  |  |  | Control | 0.267 | 0.120 | -0.050 | 50.7 | -2.22 | 3.040 | [-0.078, -0.973] |
| Valley | FAKI14 | 0.400 | 0.124 | Sucrose | 0.117 | 0.051 | 0.283 | 17.5 | 8.71 | 1.679 | [2.709, 4.795] |
|  |  |  |  | Control | 0.361 | 0.077 | 0.040 | 20.4 | 1.177 | 2.529 | - |
| Flat | FAKI14 | 0.280 | 0.073 | Sucrose | 0.115 | 0065 | 0.165 | 43.0 | 7.970 | 1.051 | [1.718, 3.374] |
|  |  |  |  | Control | 0.273 | 0.066 | 0.006 | 38.6 | 0.292 | 7.715 | - |
| FAKI14 | Valley | 0.400 | 0.124 | Hill | 0.217 | 0.051 | 0.183 | 18.3 | 5.603 | 2.393 | [1.467, 3.190] |
