## Supplementary File 8 for "Surface curvature and basal hydraulic stress induce spatial bias in cell extrusion"

| **a** Statistics of Expression Ratio of p-FAK / FAK at Various Hydraulic Stress Conditions | | | | | | | | | | | |
| --- | --- | --- | --- | --- | --- | --- | --- | --- | --- | --- | --- |
| **Conditions** | **Descriptive Statistics** | | | **One-Sample t-Test (vs. Control)** | | | | | **Cohen’s D** | | |
|  | **N** | **Mean** | **Std Dev** | **t** | **p** | **df** | **95% CI** | | **d** | **95% CI** | |
|  |  |  |  |  |  |  | **Lower** | **Upper** |  | **Lower** | **Upper** |
| **120mM Sucrose** | 5 | 126.11 | 19.57 | 2.984 | 0.0406 | 4 | 1.81 | 50.40 | 1.33 | 0.05 | 2.55 |
| **25% H_2_O** | 5 | 77.71 | 15.49 | 3.218 | 0.0324 | 4 | -41.50 | -3.06 | -1.44 | -2.70 | -0.11 |
| **CY52** | 5 | 125.55 | 53.61 | 1.024 | 0.3637 | 4 | -42.01 | 91.11 | 0.48 | -0.48 | 1.38 |
| **PAM** | 5 | 177.38 | 36.28 | 4.770 | 0.0088 | 4 | 32.34 | 122.40 | 2.13 | 0.44 | 3.78 |

| **b** Statistics of Expression Ratio of p-Akt / Akt at Various Hydraulic Stress Conditions | | | | | | | | | | | |
| --- | --- | --- | --- | --- | --- | --- | --- | --- | --- | --- | --- |
| **Conditions** | **Statistics** | | | **One-Sample t-Test (vs. Control)** | | | | | **Cohen’s D** | | |
|  | **N** | **Mean** | **Std Dev** | **t** | **p** | **df** | **95% CI** | | **d** | **95% CI** | |
|  |  |  |  |  |  |  | **Lower** | **Upper** |  | **Lower** | **Upper** |
| **120mM Sucrose** | 5 | 269.77 | 86.10 | 4.409 | 0.0116 | 4 | 62.86 | 276.70 | 1.97 | 0.37 | 3.52 |
| **25% H_2_O** | 5 | 92.08 | 17.67 | 1.002 | 0.3729 | 4 | -29.85 | 14.02 | -0.45 | -1.35 | 0.50 |
| **CY52** | 5 | 136.56 | 24.19 | 3.380 | 0.0278 | 4 | 6.53 | 66.59 | 1.47 | 0.12 | 2.75 |
| **PAM** | 5 | 286.50 | 85.78 | 4.861 | 0.0083 | 4 | 79.99 | 293.00 | 2.17 | 0.46 | 3.84 |

| **Target** | **Host** | **Type** | **Dilution** | **Incubation Time (h)** | **Manufacturer** | **Cat. No.** |
| --- | --- | --- | --- | --- | --- | --- |
| AKT (pan) | Rabbit | Monoclonal | 1:1000 | 24 | CST | 4691 |
| Phospho-AKT (Ser473) | Rabbit | Monoclonal | 1:1000 | 48 | CST | 4060 |
| FAK | Mouse | Monoclonal | 1:500 | 48 | BD Biosciences | 611008 |
| Phospho-FAK (Try397) | Rabbit | Monoclonal | 1:1000 | 48 | Thermo Scientific | 700255 |
| GAPDH | Mouse | Monoclonal | 1:1000 | 24 | Santa-Cruz Biotech | 32233 |

**c** Dilution factors, incubation duration and manufacturer’s catalogue numbers for immunoblotting.
