## Supplementary File 9 for "Surface curvature and basal hydraulic stress induce spatial bias in cell extrusion"

|  | Statistics | | | Two-tailed student’s t-test | | | | | Cohen D | | |
| --- | --- | --- | --- | --- | --- | --- | --- | --- | --- | --- | --- |
| Curvature type | N | Mean  (pFAK/FAK ratio) | SD  (pFAK/FAK ratio) | t | P | df | 95 % CI | | d | 95 % CI | |
|  |  |  |  |  |  |  | lower | upper |  | lower | upper |
| Hill | 9 | 0.882 | 0.039 | 8.330 | 3.269×10^-7^ | 16 | 0.108 | 0.182 | 3.927  (large) | 2.217 | 5.637 |
| Valley | 9 | 0.737 | 0.034 |  |  |  |  |  |  |  |  |

Statistics of pFAK/FAK fluorescence intensity ratios over hills and valleys.
